# Osteoblast-osteoclast co-cultures: a systematic review and map of available literature

**DOI:** 10.1101/2021.09.09.459671

**Authors:** S. J. A. Remmers, B. W. M. de Wildt, M. A. M. Vis, E. S. R. Spaander, R.B.M. de Vries, K. Ito, S. Hofmann

**Affiliations:** Orthopaedic Biomechanics, Department of Biomedical Engineering and the Institute of Complex Molecular Systems, Eindhoven University of Technology, Eindhoven, the Netherlands; SYRCLE, Department for Health Evidence, Radboud Institute for Health Sciences, Radboudumc, Nijmegen, the Netherlands

## Abstract

Drug research with animal models is expensive, time-consuming and translation to clinical trials is often poor, resulting in a desire to replace, reduce, and refine the use of animal models. One approach to replace and reduce the use of animal models in research is using *in vitro* cell-culture models.

To study bone physiology, bone diseases and drugs, many studies have been published using osteoblast-osteoclast co-cultures. The use of osteoblast-osteoclast co-cultures is usually not clearly mentioned in the title and abstract, making it difficult to identify these studies without a systematic search and thorough review. As a result, researchers are all developing their own methods from the ground up, leading to conceptually similar studies with many methodological differences and, as a direct consequence, incomparable results.

The aim of this study was to systematically review existing osteoblast-osteoclast co-culture studies published up to 6 January 2020, and to give an overview of their methods, predetermined outcome measures (formation and resorption, and ALP and TRAP quantification as surrogate markers for formation and resorption, respectively), and other useful parameters for analysis. Information regarding these outcome measures was extracted and collected in a database, and each study was further evaluated on whether both the osteoblasts and osteoclasts were analyzed using relevant outcome measures. From these studies, additional details on methods, cells and culture conditions were extracted into a second database to allow searching on more characteristics.

The two databases presented in this publication provide an unprecedented amount of information on cells, culture conditions and analytical techniques for using and studying osteoblast-osteoclast cocultures. They allow researchers to identify publications relevant to their specific needs and allow easy validation and comparison with existing literature. Finally, we provide the information and tools necessary for others to use, manipulate and expand the databases for their needs.

## Introduction

Bone is a highly dynamic tissue with mechanical and metabolic functions that are maintained by the process of bone remodeling by the bone forming osteoblasts (OBs), bone resorbing osteoclasts (OCs), and regulating osteocytes. In healthy tissue, bone resorption and formation are in equilibrium, maintaining the necessary bone strength and structure to meet the needs of the body. In diseases such as osteoporosis and osteopetrosis this equilibrium is disturbed, leading to pathological changes in bone mass that adversely affect the bone’s mechanical functionality (1).

Studies on bone physiology, bone disease and drug development are routinely performed in animal models, which are considered a fundamental part of preclinical research. The use of animals raises ethical concerns and is generally more time consuming and more expensive than *in vitro* research. Laboratory animals are also physiologically different from humans and their use in pre-clinical studies leads to poor translation of results to human clinical trials (2,3), and the subsequent failure of promising discoveries to enter routine clinical use (4,5). These limitations and the desire to reduce, refine and replace animal experiments gave rise to the development of *in vitro* models (6,7). Over the last four decades, significant incremental progress has been made towards developing OB-OC co-culture models.

The development of *in vitro* OB-OC co-cultures started with a publication of T.J. Chambers in 1982 (8), where the author induced quiescence of isolated tartrate resistant acid phosphatase (TRAP)-positive rat OCs with calcitonin and reversed their quiescence by co-culturing them with isolated rat OBs in direct contact. At that time, studies involving OCs resorted to the isolation of mature OCs by disaggregation from fragmented animal bones. The first account of *in vitro* osteoclastogenesis in coculture was realized in 1988 when Takahashi and co-authors (9) cultured mouse spleen cells and isolated mouse OBs in the presence of 1α,25-dihydroxyvitamin D3 and found TRAP-positive dentineresorbing cells. The herein described methods were used and adapted to generate OCs for the following decade. Most of the studies published until this point in time used co-cultures as a tool for achieving osteoclastogenesis, as opposed to a model for bone remodeling. At that time, a co-culture of OBs with spleen cells or monocytes was the only way of generating functional OCs *in vitro.* It wasn’t until 1999 that Suda (10) discovered Receptor Activator of Nuclear Factor Kappa Ligand (RANKL) and Macrophage Colony Stimulating Factor (M-CSF) as the necessary and sufficient proteins required for differentiating cells from the monocyte/macrophage lineage into functioning OCs (11–13). This discovery marked the start of co-culture models developed for studying bone remodeling.

In recent years, many research groups have ventured into the realm of OB-OC co-cultures with the intent of studying both formation and resorption, but each group seems to be individually developing the tools to suit their needs resulting in many functionally related experiments that are methodologically completely different. In addition, the use of such methods is often not clearly stated within title and abstracts. Simple title/abstract searches such as ‘OB + OC +co-culture’ tend to scratch only the surface of the base of evidence available using OB-OC co-cultures. Finding and comparing different co-culture approaches and results is thus virtually impossible and forces each group to develop and use their own methods instead of building upon those of others.

The aim of this study was to construct a systematic review of all OB-OC co-cultures published up to January 6, 2020. With this systematic review, we aimed at identifying all existing OB-OC co-culture studies and analyze these within two comprehensive databases, allowing researchers to quickly search, sort and select studies relevant for their own research. Database 1 contains all OB-OC coculture studies in which at least one relevant primary outcome measure was investigated (formation and/or resorption) or secondary outcome measure (alkaline phosphatase (ALP) and/or tartrate resistant acid phosphatase (TRAP) quantification as surrogate markers for formation and resorption, respectively) (S1_File_Database_1). A sub-selection of studies that investigated these relevant outcome measures on both OBs and OCs in the co-culture was included in Database 2, accompanied by additional details on methods, culture conditions and cells (S2_File_Database_2). The collection of the two databases will further be referred to as a systematic map.

## Methods

For this systematic map a structured search protocol was developed using the SYRCLE protocol format (14). This protocol format is tailored to the preparation, registration, and publication of systematic reviews of preclinical studies, and helps authors predefine the methodological approach of their review from research question to data synthesis. The protocol and search strings were made publicly available before completion of the study selection via Zenodo (15) to ensure transparency of the publication. In short, three online bibliographic literature sources were consulted with a comprehensive search string and the resulting publications were combined and screened using a four-step procedure (Fig. 1): 1) identification of OB-OC co-cultures, 2) identification of relevant outcome measures, 3) categorization in Databases 1 and 2 (Fig 2), 4) search for additional articles in the reference lists of studies included in Database 2 and relevant reviews.

**Fig 1.**
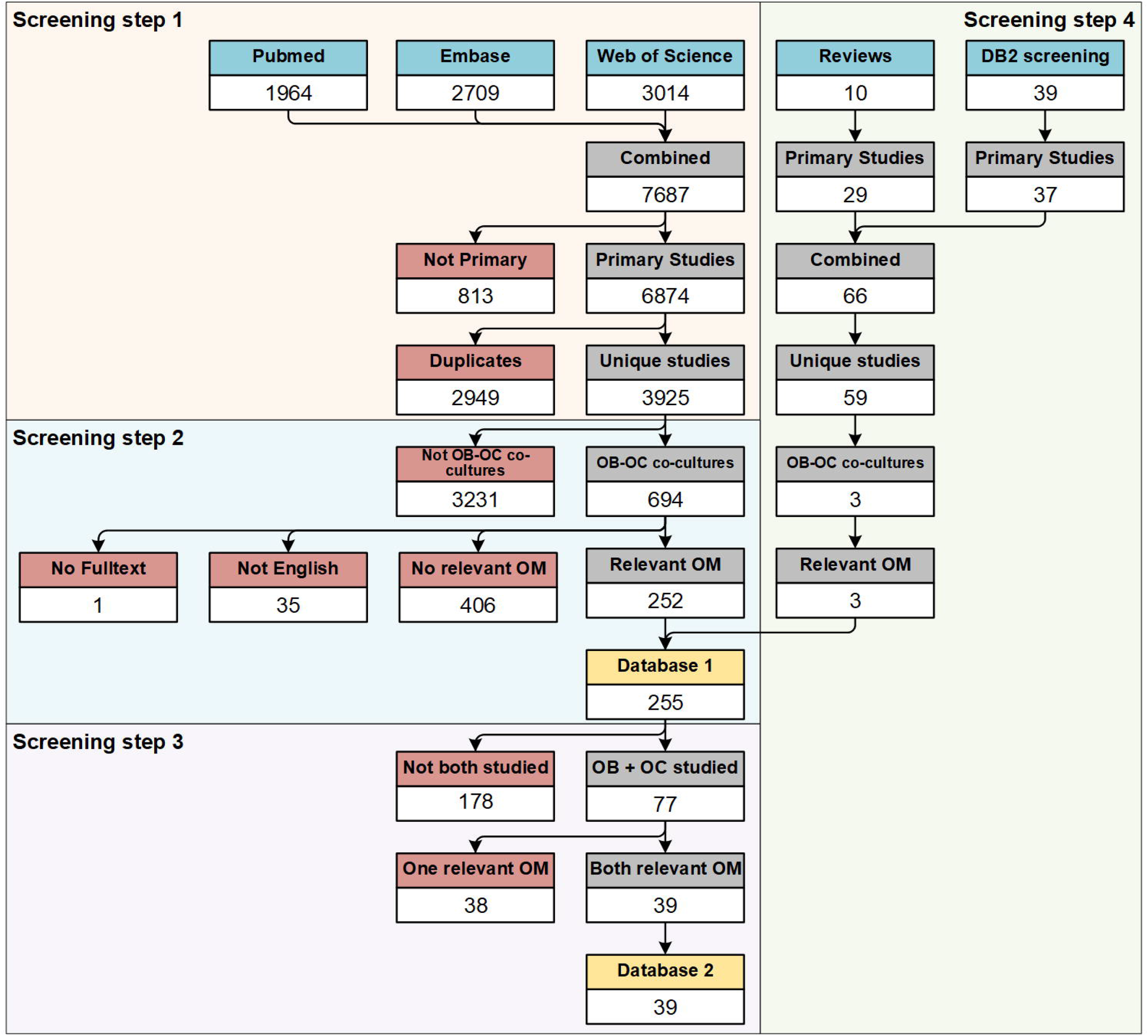
Flow diagram of systematic literature search and screening. Screening step 1: Hits from 3 online bibliographic literature sources were combined, primary studies were selected, and duplicates were removed. Title and abstracts were screened for the presence of OB-OC co-cultures. Screening step 2: OB-OC co-cultures were screened in full text for relevant outcome measures. All studies in which at least one relevant outcome measure was studied were included into Database 1. Screening step 3: Papers in which both cell types were studied with relevant outcome measures were included into Database 2. Screening step 4: Papers included into Database 2 and relevant reviews were screened for potentially missing relevant studies and identified studies were screened in the same manner as described here. Each screening step is marked with a separate background color. Each selection step within the screening steps is marked with a colored header. Blue header: used as input for the review. Grey header: selection step. Red header: excluded studies. Yellow header: Database as presented in this systematic map. Abbreviations: outcome measures (OM), Database 2 (DB2), osteoblast (OB), osteoclast (OC).

**Fig 2.**
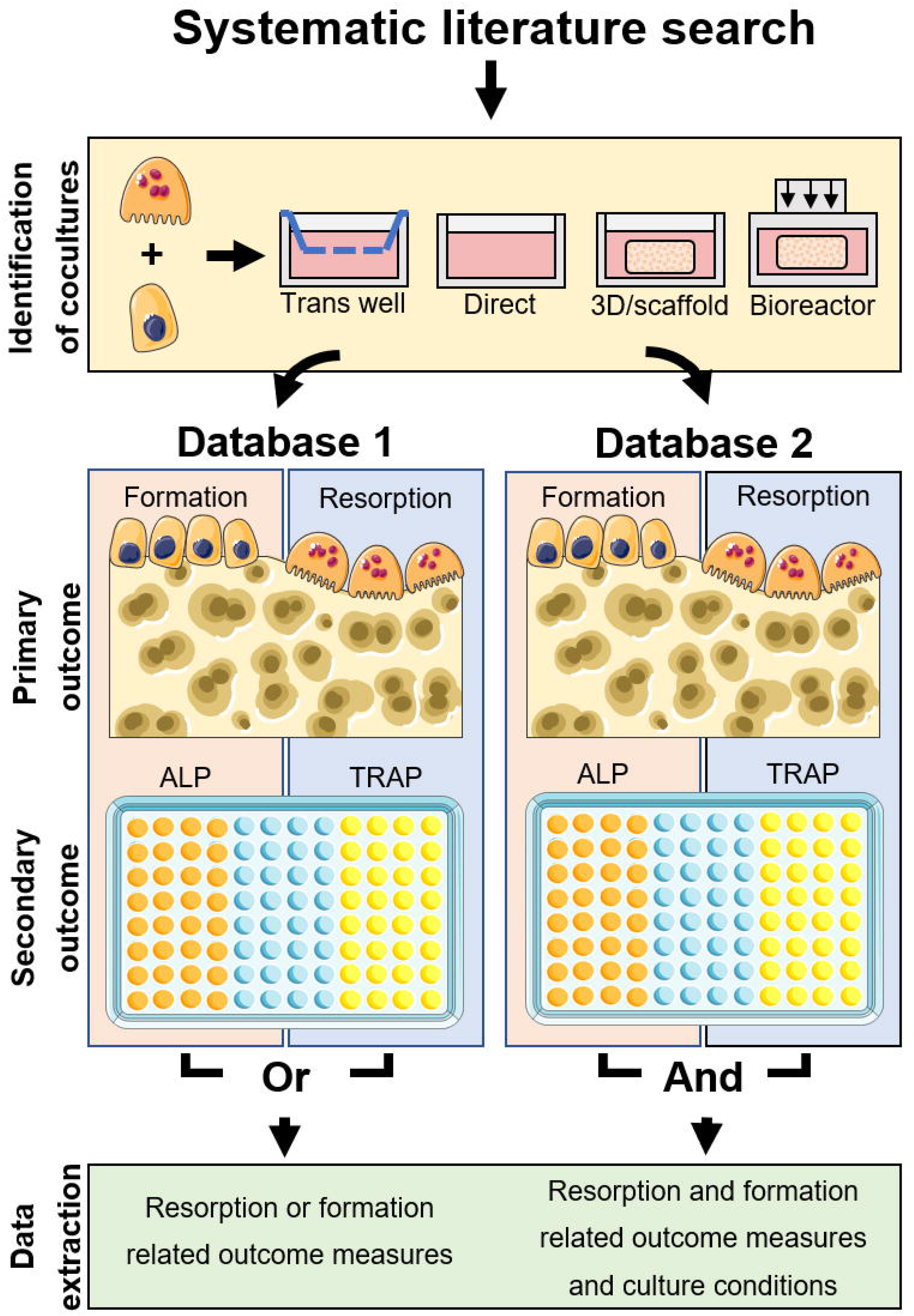
Schematic overview of Databases 1 and 2. All identified studies were searched for OB-OC cocultures, where co-culture was defined as OB and OC being present simultaneously and able to exchange biochemical signals. In addition to direct-contact cultures, cultures such as transwell cultures, 3D or scaffold cultures and bioreactor cultures were allowed as well. OB-OC co-culture studies which used relevant outcome measures were included into Database 1. Of these, only the relevant outcome measures were analyzed. All studies where relevant outcome measures were used for both OB and OC were included into Database 2 as well. Of these, cells and culture conditions were analyzed. The figure was modified from Servier Medical Art, licensed under a Creative Common Attribution 3.0 Generic License (http://smart.servier.com, accessed on 2 July 2021).

### Database Search

The online bibliographic literature sources Pubmed, Embase (via OvidSP) and Web of Science were searched on January 6, 2020 with a predefined search query developed to identify as many studies as possible employing OB-OC co-cultures. The search strings used a combination of thesaurus and free text terms where possible and consisted of the following components: ([OBs] OR ([OB precursors] AND [bone-related terms])) AND ([OCs] OR ([OC precursors] AND [bone-related terms])) AND [co-culture], where each component in square brackets represents a list of related thesaurus and free-text search terms, and where parentheses indicate the order of operations within the search query. The full search strings can be found via Zenodo (15). The results of all three searches were combined. Conference abstracts and duplicates were removed using the duplicate removal tools of Endnote X7 and Rayyan web-based systematic review software (16).

The remaining entries were screened using a four-step procedure that resulted in the generation of 2 databases: Database 1 containing all co-culture studies that measured at least one relevant outcome measure (formation, resorption, ALP or TRAP), and Database 2 containing all studies that measured at least one relevant outcome measure on both OB and OC: either formation or ALP for OB, and either resorption or TRAP for OC. The entire data collection and screening process was performed independently by two researchers starting from the moment that conference abstracts and duplicates had been removed and ending the moment that the relevant information of the publications was extracted into the two databases. Disagreements between researchers were discussed and publications were re-screened if necessary. A third independent researcher was consulted for disagreements that could not be solved by the first two independent researchers.

#### Screening step 1: Identification of OB-OC co-cultures

This step was performed to identify and extract OB-OC co-cultures from the complete list of studies identified from the three online bibliographic literature sources after automatic removal of conference abstracts and duplicates. All further steps were done on these studies or a sub-selection thereof. Using Rayyan web-based systematic review software (16), the titles and abstracts of all entries were screened for the presence of primary studies using OB-OC co-cultures. Reviews, theses, chapters, and conference abstracts that were not automatically detected were excluded at this point. Potentially relevant reviews were saved separately to serve as an additional source of studies that could have been missed by the systematic search.

In the selection process, co-culture was defined as the simultaneous presence (verified) or assumed presence (expected) of OBs and OCs (or OB-like and/or OC-like cells) within the same culture system at a moment during the described experiment such that the cells were able to communicate either via soluble factors in the medium and/or direct cell-cell contact. Both primary cells and cell lines of any origin were admitted including heterogeneous cell populations, if these were clearly defined and expected to result in a biologically relevant number of the desired cell type, precursor type, or terminally differentiated cell type. The presence of progenitor cells (such as monocytes or mesenchymal stem/stromal cells) was allowed only if these were either verified or expected to differentiate into OBs and/or OCs. Studies using a single animal or human donor for both cell types were allowed, but only if the two (progenitor) cell types were at one point separated, counted, and reintroduced in a controlled manner. In addition, trans-well systems (no physical contact but shared medium compartment with or without membrane), scaffolds (3-dimensional porous structure of any material including decellularized matrix), and bioreactor culture systems (culture exposed to physical stimuli such as rotation, mechanical loading or fluid flow) were included. Conditioned media experiments were excluded because these do not allow real-time two-way exchange of cell signals. Explant cultures or organ cultures were excluded because these studies contain a living ex *vivo* culture element, whereas the focus of this systematic map is limited to *in vitro* studies.

When the study used any type of OB-OC co-culture as defined above, the study was included. When there was no indication that there was an OB-OC co-culture, the study was excluded. When, based on the title and abstract, it was likely that there was a co-culture, but this was not described as such, the full-text publication was screened.

#### Screening step 2: Identification of relevant outcome measures in the coculture experiments

This step was used to identify co-cultures that specifically investigated outcome measures related to bone remodeling: measuring formation or resorption (primary outcome measures), or quantitative measurements of activity markers ALP or TRAP in a dedicated assay (secondary outcome measures). The primary outcome measures of measuring resorption and formation were chosen because these are the processes that are directly affected in bone diseases. Measuring these outcome measures usually requires a specific methodological setup such as a specific surface analysis for measuring formation, or a resorbable substrate for measuring resorption. The secondary outcome measures of ALP and TRAP were included because these are regarded as viable alternatives for the direct measurement of formation and resorption. The full texts of the studies identified in screening step 1 were screened for experimental techniques and outcome measures. Studies in which for at least one of the cell types a relevant outcome measure was used, were selected to be used in Database 1 (S1_File_Database_1). The measurement of formation was defined as any method that directly measures the area or volume of (tissue) mineralization by OBs, any method that measures byproducts of formation, and any method that measures biochemical markers that directly and exclusively correlate to formation. The measurement of resorption was defined as any method that directly measures the surface area or volume that has been resorbed by OCs, that biochemically measures products or by-products of resorption, or that measures biochemical markers that directly and exclusively correlate to resorption. The measurement of ALP and TRAP was defined as the detection of either the direct measurement of the enzymatic activity of these proteins, or the direct quantification of the amount of those proteins present in a dedicated assay. Studies that determined ALP or TRAP gene expression using PCR were excluded because PCR was not considered a dedicated assay for this map and did not directly measure the amount of protein present. However, the use of PCR was recorded in the generated databases in a separate column. Immuno-histological stainings of ALP or TRAP were not considered relevant outcome measures, even when followed by image analysis because at best these quantify stained surface area and not actual protein content.

All co-cultures that did not contain at least a single outcome measure that met these criteria were excluded from further use. Because this was the first step at which the availability of the full text publication was required, publications written in languages other than English with no translation available, and publications of which the full text could not be found were excluded at this point.

#### Screening step 3: Categorization within Database 1

In this step, a distinction was made between studies in which only one of either OB or OC was studied, or both were studied. This distinction was made because ideally, a model for bone remodeling should show effects on both OBs and OCs. Each study selected from screening step 2 was assessed on the methods to study OBs and OCs. Each study was categorized into one of five categories within Database 1: 1) The relevant outcome measures were measured in both OBs and OCs in the co-culture. These studies were also included in the in-depth screening for Database 2 (S2_File_Database_2). 2) Both cell types were studied, but relevant outcome measures were only measured in OCs or 3) Both cell types were studied, but relevant outcome measures were only measured in OBs. 4) Only OCs were studied in co-culture, the other cell type was neglected or 5) OBs were studied in co-culture, the other cell type was neglected. Thus, category 1 contained the studies in which both formation and resorption were investigated, either directly or by ALP or TRAP quantification. Category 2 and 3 contained studies in which both OBs and OCs were studied, but only one of the two was studied with the relevant outcome measures. The other cell type was studied using other methods instead such as stainings or PCR. Categories 4 and 5 contain studies in which only one of the two cell types was analyzed with one of the relevant outcome measures while the other cell type was present but not analyzed in co-culture at all. Note that for this categorization, it was necessary that the cells that were used in co-culture were studied, and not for example a monoculture conducted in parallel.

#### Screening step 4: Review and reference list screening

To find additional studies that may have been missed during bibliographic searches, relevant review articles identified during the selection process and studies labeled as category 1 in step 3 were screened for additional publications that could be relevant to the current systematic map. Of these studies, the relevant passages within the text were screened, followed by a thorough screening of the complete reference lists of these studies. All potentially relevant studies were first cross-checked with the original search results of the bibliographic literature search, and if these were not identified there, were screened in the same manner as all other studies used in this systematic map. Unique relevant studies were then added to the corresponding databases and analyzed as described earlier.

### Database 1 generation and analysis – All co-cultures with relevant outcome measures

Every study included in Database 1 was screened for the relevant outcome measures resorption, formation, ALP and TRAP during screening steps 2 and 3. To provide useful and specific information of each of the studies included in this database, all potentially relevant information related to the relevant outcome measures was collected and organized. For resorption, additional information on the resorbed substrate, the methodological procedure and quantification of results was collected. For formation, additional information on the type of analysis, the methodological procedure and quantification of results was collected. For both ALP and TRAP, additional information on the mechanism of the biochemical assay, whether it was conducted on lysed cells or supernatant, and information regarding the quantification was collected. In addition, the following information was extracted, whether: the authors described their setup as a model specifically for remodeling, the experiment was conducted in 3D, the experiment applied bioreactors, more than 2 cell types were cultured simultaneously, the culture used a trans-well setup, the culture used PCR and components in the supernatant of the culture were analyzed by ELISA or a similar quantification method. If the answer to these questions was yes, then the applicable details were collected as well. Finally, a column for additional remarks was introduced for details that did not fit in another column. Studies where the authors are color coded in pink were those not found through the initial database search but by the screening of the review articles or reference lists. Studies categorized as category 1 in screening step 3 were selected for use in Database 2 and had their title color coded in orange.

#### Quality assessment and scripting

In Database 1 only the methods used for analyzing relevant outcome measured are reported, and not the data obtained from them or the results described in the publication. Quality assessment in Database 1 is thus limited to assessing the completeness of the necessary elements of the collected methodological details, to the extent that the methods are properly represented in Database 1 and related tables. Please note that the methods themselves were not investigated on a complete description for a perfect replication of the study, but only on the description necessary to accurately classify the method within this systematic map. For example, a study claiming to investigate resorption on dentine discs using Toluidine Blue was deemed sufficiently described to accurately classify, regardless of whether the information presented was sufficient to duplicate that specific method precisely. Publications in which information was missing are here represented as ‘not reported’ if no information was provided, ‘reference only’ if no information was provided but another study was referenced, and ‘undefined kit’, when a commercial kit was used but the content or methodology was not further described. Instances of missing information can easily be identified in figures, tables and databases, but were not further used in this systematic map. Studies where an instance of information was missing were still used for other analyses for which the corresponding provided information was present.

A script was written in Excel Visual Basics programming language to analyze Database 1 and extract relevant statistical information on the collected information. On sheet 2 “Data” of the Database 1 excel file, the descriptive statistical data and collected information are presented in the form of lists and tables and together with a button to re-run the analysis based on the reader’s requirements. The script is integrated within the excel file and can be used only when the file is saved as a ‘macro-enabled’ file (.xlsm).

### Database 2 generation and analysis – All co-cultures in which both cell types had relevant outcome measures

In addition to the information already collected for Database 1, additional information was extracted from the studies in which relevant outcome measures were studied of both OB and OC: the species (17) and type (cell line or primary) and actual used cell type (6) of both the OB and OC were collected. Seeding numbers and densities (18) for both OB and OC were collected or calculated where possible, separated by 2D (cell density per area) and 3D (cell density per volume), and the seeding ratio (19) between OB and OC was noted or calculated. The culture surface (bio-)material (20), sample size (samples per group), culture duration and medium refreshing rate in units as reported in the study, environmental conditions or variations such as CO_2_ or O_2_ alterations or mechanical loading (21), and pre-culture duration (22) were collected, where pre-culture is defined as a different co-culture condition (such as a different supplement cocktail) lasting for a short duration (such as 2 days) prior to the ‘main’ co-culture. The medium composition (23) was collected and organized by base medium type such as Dulbecco’s Modified Eagle Medium (DMEM) and alpha-Modified Eagle Medium (αMEM), glucose content (if provided separately in the text), Fetal Bovine Serum (FBS) / Fetal Calf Serum (FCS) in percentages, antibiotics (types and concentrations or percentages as provided), OB supplement concentrations (ascorbic acid, β-glycerophosphate and dexamethasone, OC supplement concentrations (M-CSF and RANKL) and other supplements, as well as medium content of any monoculture prior to the co-culture. Finally, the tested genes of all studies applying PCR and any proteins studied with ELISA or other supernatant analyses executed on the coculture were noted.

#### Quality assessment and scripting

In Database 2 the culture conditions, cells and materials used are reported, and not the data obtained from them or the results described in the publication. Quality assessment in Database 2 is thus limited to assessing the completeness of the necessary elements of the collected methodological details, to the extent that the methods are properly represented in Database 2 and related figures and tables. Please note that the methods themselves were not investigated on a complete description for a perfect replication of the study, but only on the description necessary to accurately classify the method within this systematic map. For example, a study claiming to use human primary monocytes and human primary osteoblasts for the OB-OC co-culture was deemed sufficiently described to accurately classify respectively the OB and OC origin, regardless of whether the information presented was sufficient to perfectly replicate that part of the study. Publications in which information was missing are here represented as ‘not reported’ (NR) if no information was provided, or ‘reference only’ if no information was provided but another study was referenced. If studies were missing information critical to reproduce the outcome measures (for example seeding ratio’s, culture surface material, medium or supplement information, critical steps in analyses), the cells in the database missing this information were labeled in red. If the missing information was not critical for the outcome measures but necessary for a replication of the study (for example sample size, medium refresh rate, control conditions), the cells were labeled in orange. The sum of both orange and red cells for each color in each study is shown as well to indicate how many instances of missing information were identified in each study. The color coding was determined by the authors of this map but can be adjusted within Database 2 if other criteria for critical information and completeness are desired. Instances of missing information can easily be identified in the corresponding figures, tables and databases, but were not further used in this systematic map. Studies where an instance of information was missing were still used for other analyses for which the corresponding provided information was present.

Using Excel Visual Basics programming language, three scripts were written to analyze and process Database 2. One script was created to count all instances of cells labeled as ‘missing info’ and present this number in two dedicated columns (missing critical or non-critical info). One script was created to count the frequency of occurrence of all (co-)authors and years of publication. Finally, one script was created to analyze this database and extract relevant descriptive statistical data on the collected information. On sheet 2 “Data” of the Database 2 excel file, the statistical data and collected information are presented in the form of lists and tables and together with the buttons to re-run the analyses based on the reader’s requirements. The scripts are integrated within the excel file and can be used only when the file is saved as a ‘macro-enabled’ file (.xlsm).

## Results

### Search results

From three online bibliographic literature sources, 7687 studies were identified (Pubmed: 1964, Embase via OvidSP: 2709, Web of Science: 3014). After removal of conference abstracts, 6874 studies remained. After duplicate removal, 3925 unique studies were identified to be screened.

#### Studies included into Database 1

After title-abstract screening and when in doubt full text screening (screening step 1), 694 studies were identified as OB-OC co-cultures. A list of these studies is available as a supplementary file (S4_File_List of all OB-OC co-cultures). Of these, one study was excluded from further analysis because the full text could not be obtained, 35 were excluded because they were in a language other than English and 406 were excluded because no relevant outcome measure was used in the study (screening step 2). The qualifying 252 studies with at least one relevant outcome measure were included in Database 1.

#### Studies included into Database 2

In 77 of these studies, both the OB and OC were studied, and in 39 of these, both OB and OC were studied using relevant outcome measures (screening step 3). These 39 studies were included in Database 2.

#### Additional screening of review articles and reference lists for missing studies

The 39 studies of Database 2 and 10 additional review publications were screened for other relevant studies that the initial search may have missed (screening step 4). An additional 25 unique studies were identified in the 10 reviews, and 34 unique studies were identified from the reference lists of the included studies. These additional 59 studies were reviewed as described previously and resulted in an additional 3 OB-OC co-cultures with only relevant outcome measures measured on one cell type, resulting in a total of 255 studies with relevant outcome measures on at least one cell type for Database 1, and 39 studies in which relevant outcome measures were studied in both cell types for Database 2. A detailed overview of the search and selection process is shown in Fig 1.

#### Publications per year

The publications included in Database 1 were published between 1983 and 2019, with a peak in publications around the year 2000, followed by a dip and then a more or less slight but steady increase until now (Fig 3a). The peak roughly coincides with the discovery that M-CSF and RANKL were both necessary and sufficient to induce osteoclastic differentiation in monocytes in 1999 (10). The included publications in Database 2 span the time between 1997 and 2019, with only 8 publications before 2010 (Fig 3b). This coincides with the progress in development of *in vitro* cocultures of OB and OC, moving beyond co-cultures with OB to generate OC, and moving towards cocultures of OB and OC to study for example cell-cell interactions (6).

**Fig 3.**
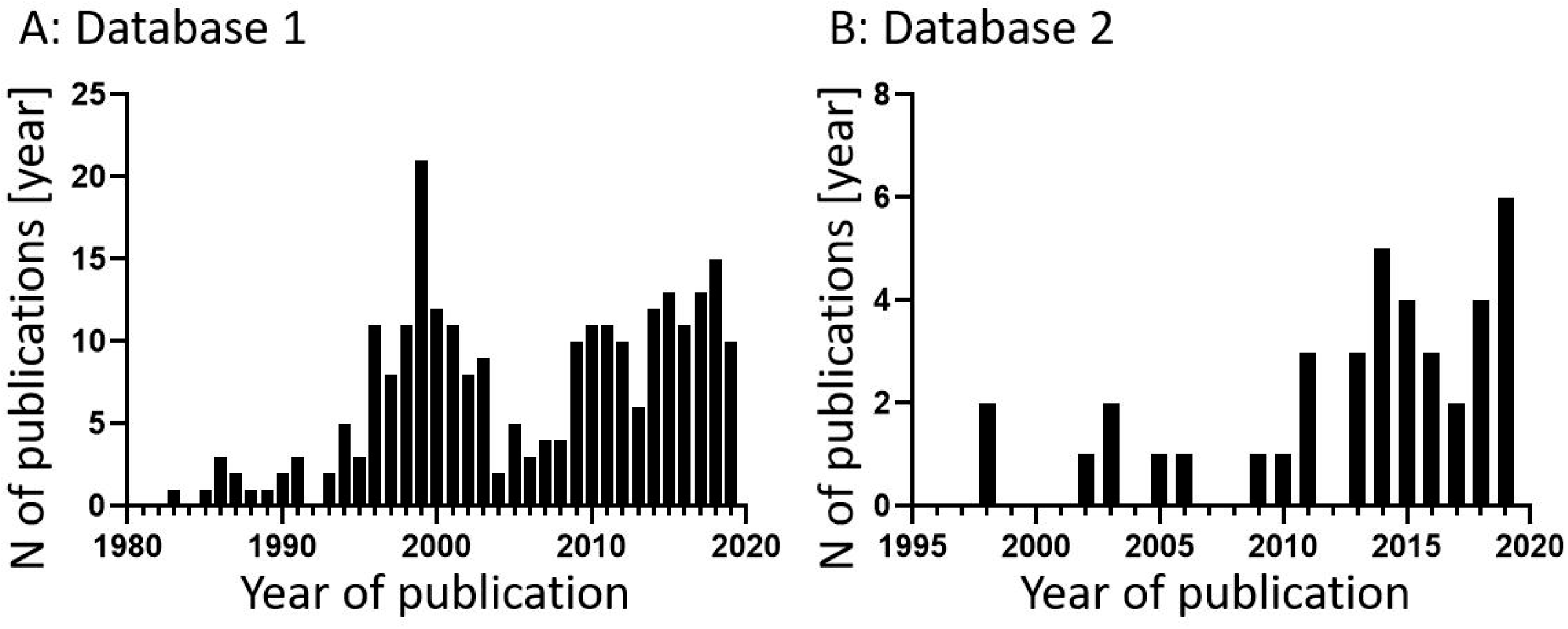
Relevant publications per year. A) All 255 publications that contain relevant outcome measures counted by year ranging from 1983 to 2019 (Database 1). B) The 39 selected publications of Database 2 counted by year ranging from 1998 to 2019 (Database 2).

### Database 1 results

Database 1 provides an overview of all OB-OC co-culture studies published until January 6, 2020 in which at least one relevant outcome measure was studied. Of the 255 studies included, resorption was analyzed in 181 studies, formation was analyzed in 37 studies and both were analyzed in 16 studies. ALP was analyzed in 42 studies, TRAP was analyzed in 61 studies and both were analyzed in 22 studies (Table 1).

**Table 1.** Combinations and frequencies of primary and secondary outcome measures. This table can be referenced to identify the number of studies using any combination of primary and secondary outcome measures. All 255 studies that investigate at least one of the primary or secondary outcome measures are represented once in this table. Each study is represented by a combination of primary outcome measures (horizontal) and secondary outcome measures (vertical). Marginal totals of each row and column are counted under ‘total’ with the grand total in the bottom-right cell. These marginal totals sum the total number of studies that studied only that combination of either primary or secondary outcome measures, with no regard of the outcome measures on the opposite axis. The total numbers of each individual outcome measure can be calculated from this table but are presented in the following paragraphs and tables.

| Combinations of primary and secondary outcome measures in each study |  | Primary outcome measures |  |  |  |  |
| --- | --- | --- | --- | --- | --- | --- |
|  |  | No resorption or formation | Resorption only | Formation only | Resorption and formation | Total |
| Secondary > | No ALP or TRAP | 0 | 151 | 14 | 9 | 174 |
|  | ALP only | 16 | 0 | 2 | 2 | 20 |
|  | TRAP only | 23 | 9 | 3 | 4 | 39 |
|  | ALP + TRAP | 14 | 5 | 2 | 1 | 22 |
|  | Total | 53 | 165 | 21 | 16 | 255 |

#### Resorption

Resorption is the process by which osteoclasts remove old and damaged bone tissue through enzymatic degradation or acidic dissolution. Out of all 255 OB-OC co-culture publications included in Database 1, resorption was studied directly on 188 occasions in 181 studies and quantified 142 times (Table 2). In some publications, more than one material or method of analysis for resorption was used. In cases where multiple materials were used, each material was counted as an individual study of resorption and for each material, the corresponding analyses were counted, even if these were identical per material. In those cases where multiple methods of analysis are used on the same material, all methods are counted, and the material is counted only once. This resulted in a counted number of studies that is higher than the actual number of publications. When numbers of studies are referenced, these are the ‘counted’ number of studies defined above, and not the actual number of publications.

**Table 2a.** Occurrences of resorption on different types of substrates and subsequent analyses. Each column signifies a different material used as a substrate for measuring resorption. If other cells, prior to the introduction of OC, were used to deposit mineralized matrix on another material, then the material was listed in the column ‘Mineralized’. If the material was not reported, the study was listed in the column ‘Not reported’. The first rows show how many instances of each material were included into this systematic map in total, and how many times the results were quantified. The final column shows incremental totals per material type or analysis type. This table consists of two sections. The top section shows in what form or shape the corresponding materials were used as a substrate for resorption. The bottom section shows the techniques that were used to study the resorption described on the materials described in the top section. Each individual study is represented exactly once in the top section of the table to signify the type and form of the substrate used, and exactly once in the bottom section of the table to signify the method used to analyze the resorption that occurred on that substrate. This required the selection of the most ‘important’ part of the methods used. In the cases where first a staining was used followed by microscopy, only the staining is listed. Only in those cases where resorption was investigated directly with a microscope without prior staining, the type of microscopy is listed.

|  |  | Materials used as a resorbable substrate for measuring resorption |  |  |  |  |  |  |  |  |  |  |  |
| --- | --- | --- | --- | --- | --- | --- | --- | --- | --- | --- | --- | --- | --- |
| Shapes, structures and types of materials used as resorbable substrate for analysis of resorption. |  | Dentine | Bone | HA | Silk | Collagen | CaP | PLLA | Chitosan | Osteologic | Mineralized | Not reported | Per-row total |
| Per-material total number of studies |  | 76 | 66 | 6 | 5 | 2 | 4 | 1 | 1 | 19 | 6 | 2 | 188 |
| Per material quantified studies |  | 55 | 52 | 3 | 4 | 1 | 4 | 0 | 0 | 17 | 4 | 2 | 142 |
| Shape or structure of material | Discs | 76 | 63 | 2 |  |  | 2 |  |  | 13 |  |  | 156 |
|  | Films |  |  | 2 | 4 |  |  | 1 | 1 |  |  |  | 8 |
|  | Coatings |  |  | 2 |  |  | 1 |  |  |  |  |  | 3 |
|  | Scaffolds |  |  |  | 1 | 1 |  |  |  |  | 3 |  | 5 |
|  | Hydrogels |  |  |  |  | 1 |  |  |  |  |  |  | 1 |
|  | ECM |  |  |  |  |  |  |  |  |  | 2 |  | 2 |
|  | Nodule |  |  |  |  |  |  |  |  |  | 1 |  | 1 |
|  | Fragments |  | 3 |  |  |  |  |  |  |  |  |  | 3 |
|  | Substrates |  |  |  |  |  | 1 |  |  |  |  |  | 1 |
|  | Plates |  |  |  |  |  |  |  |  | 6 |  |  | 6 |
|  | Not reported |  |  |  |  |  |  |  |  |  |  | 2 | 2 |
| Analysis techniques for analyzing resorption on resorbable substrates. |  | Dentine | Bone | HA | Silk | Collagen | CaP | PLLA | Chitosan | Osteologic | Mineralized | Not reported | Per-row total |
| Staining | Toluidine Blue | 36 | 19 |  |  |  |  |  |  |  |  |  | 55 |
|  | Haematoxylin | 16 | 2 |  |  |  |  |  |  |  |  |  | 18 |
|  | Eosin |  | 1 |  |  |  |  |  |  |  |  |  | 1 |
|  | H&E |  | 1 |  |  |  |  |  |  |  |  |  | 1 |
|  | Alum / Coomassie Blue |  | 1 |  |  |  |  |  |  |  |  |  | 1 |
|  | TRAP | 1 |  |  |  |  |  |  |  |  |  |  | 1 |
|  | Von Kossa |  |  |  |  |  | 2 |  |  | 4 | 1 |  | 7 |
| Microscopy | Phase contrast |  |  |  |  |  | 1 |  |  | 4 |  |  | 5 |
|  | SEM | 12 | 37 | 5 | 3 |  | 1 |  |  | 1 |  |  | 59 |
|  | TEM |  |  |  |  | 1 |  |  |  |  | 1 |  | 2 |
|  | 2-Photon |  |  |  |  |  |  |  |  |  | 1 |  | 1 |
|  | Atomic force |  |  |  | 1 |  |  | 1 | 1 |  |  |  | 3 |
|  | Reflected light | 8 | 2 |  |  |  |  |  |  |  |  |  | 10 |
|  | Dark field |  |  |  |  |  |  |  |  | 1 |  |  | 1 |
|  | Light microscopy |  |  |  |  |  |  |  |  | 6 |  |  | 6 |
| Other | Assay |  |  |  |  | 1 |  |  |  |  |  |  | 1 |
|  | Immuno-assay |  | 3 |  |  |  |  |  |  |  | 3 | 1 | 7 |
|  | MicroCT |  |  |  | 1 |  |  |  |  | 1 |  |  | 2 |
|  | Reference only | 2 |  |  |  |  |  |  |  |  |  |  | 2 |
|  | Not reported | 1 |  | 1 |  |  |  |  |  | 2 |  | 1 | 5 |
|  | Total per material | 76 | 66 | 6 | 5 | 2 | 4 | 1 | 1 | 19 | 6 | 2 | 188 |

**Table 2b.** Supernatant resorption techniques. This table presents five types of resorption analysis where measurements can be performed in the culture supernatant and not on the material itself. In the corresponding studies, these were done in addition to ‘regular’ analysis presented in Table 2a, and for that reason are presented separately. These have the advantage that they can be used to monitor changes over time in a non-destructive way.

| Supernatant Analysis techniques per material used for analysis of resorption. |  | dentine | bone | HA | silk | collagen | CaP | PLLA | chitosan | osteologic | mineralized | not reported | Per-row total |
| --- | --- | --- | --- | --- | --- | --- | --- | --- | --- | --- | --- | --- | --- |
| Supernatant analysis | NTx | 1 |  |  |  |  |  |  |  |  | 2 |  | 3 |
|  | CTx |  |  |  |  |  |  |  |  |  | 1 | 1 | 2 |
|  | ICTP |  | 1 |  |  |  |  |  |  |  |  |  | 1 |
|  | Phosphate release |  |  |  |  | 1 |  |  |  |  | 2 |  | 3 |
|  | Radioactive proline release |  | 2 |  |  |  |  |  |  |  |  |  | 2 |

Most of these studies used discs or fragments of either bone or dentine. Due to the flat nature of these discs, the surface can be considered 2D, and resorption pits can be visualized directly using conventional microscopy techniques, such as for example Scanning Electron Microscopy (SEM) or Reflected Light Microscopy (RLM). To enhance the contrast of the resorption pits, stains such as Toluidine Blue (TB) and Hematoxylin (H) were used. Resorption on bone fragments was quantified using radioimmunological assays measuring the release of *in vivo* pre-labeled ^3^H-proline or type I collagen telopeptide.

Synthetic resorbable discs or coatings on culture plates are designed specifically for studying resorption, and usually the exact composition has not been revealed. These will further be referred to as ‘osteologic’ plates or discs. The discs were analyzed in roughly the same way as bone or dentine discs. Thin resorbable coatings on translucent culture plates offer another interesting approach. Resorbed areas reveal the translucent culture plate, while unresorbed areas are less translucent and can be stained with for example von Kossa’s method to provide even more contrast, making quantification with image analysis easy.

Hydroxyapatite (HA) and other calcium phosphates were used in the form of discs, films, coatings, or scaffolds and were analyzed using various types of microscopy, both with and without prior staining. These can be used in a similar way as biological and synthetic materials mentioned earlier, with the main advantage being their known composition.

Resorption of ECM or nodules produced by OBs and scaffolds mineralized by OBs were investigated with transmission electron microscopy, light microscopy after staining, using 2-photon Second Harmonic Generation microscopy (24), supernatant phosphate levels, or with an ELISA for C-terminal telopeptide (CTx) or N-terminal telopeptide (NTx), which are bone turnover marker more commonly used for testing urine and serum samples.

#### Formation

Formation is the process by which osteoblasts create new bone tissue through the mineralization of deposited collagenous extracellular matrix. Out of all OB-OC co-cultures included in Database 1, formation was studied directly 39 times in 37 studies and quantified 29 times. (Table 3) In some studies, more than one method of measuring, analyzing and quantifying formation was used. In those cases, all methods are counted as individual studies. The methods of formation analysis were divided into 5 types: nodule analysis, volume analysis, surface analysis, supernatant analysis and 3D scans.

**Table 3:**
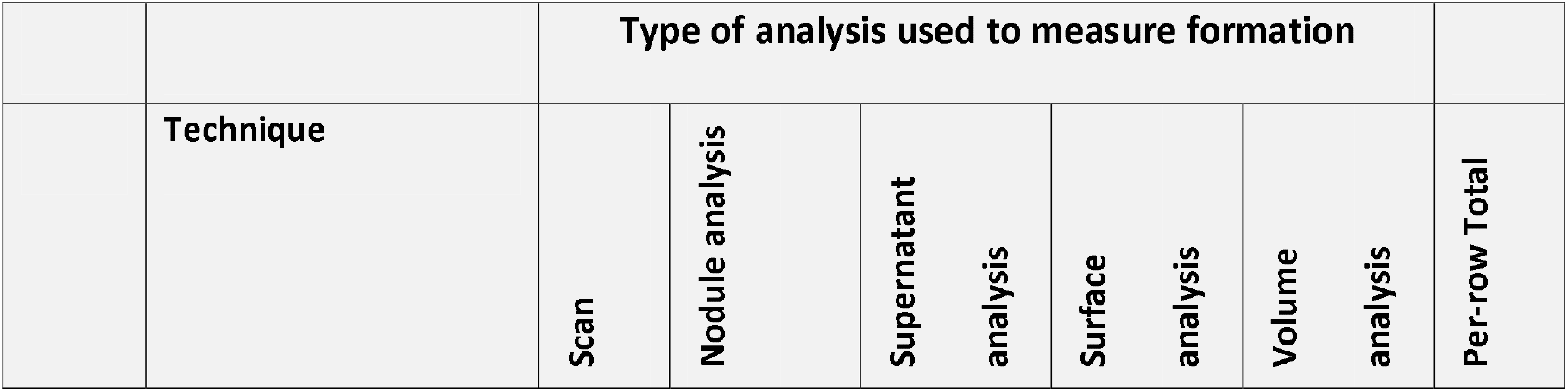
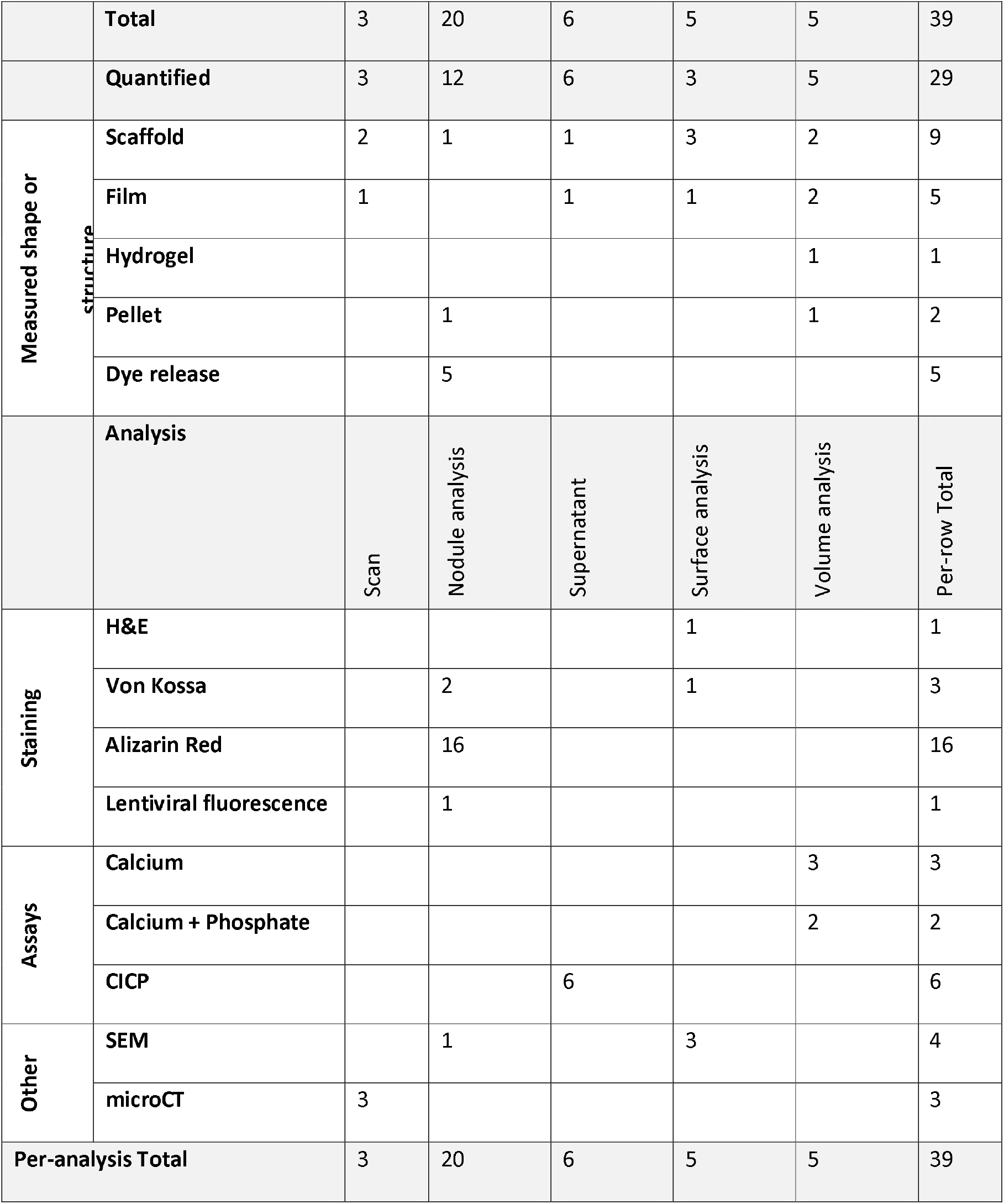
Formation statistics and analyses. Each column signifies a different type of analysis used for measuring formation. These include any type of non-destructive scan, a form of analysis of mineralized nodules, supernatant analysis, surface analysis, and destructive analysis of a mineralized volume. The first rows show how many instances of each type of analysis were included into this systematic map in total, and how many times the results were quantified. The final column shows marginal totals per row of each row. This table consists of two distinct sections, each starting with a row showing all analysis types for convenience. The first section lists defining characteristics of studies such as using films, scaffolds, hydrogels or pellets, or using a technique to first stain tissue, and then releasing and measuring the released dye. Not each study had such defining characteristics, and the total of section one does not add up to 39 studies. Section two shows either which materials was measured, or which technique was used for measuring formation. Each instance of formation is represented in section two of this table exactly once. Stainings were followed by microscopy or an assay in those cases where dye was released to be measured.

The most common method to quantify formation was to investigate mineralized nodule formation. This was done by using staining techniques such as Alizarin Red (calcium) (25) or von Kossa (phosphate) (26) followed by imaging, or directly imaging the nodules. While any staining specific for mineralized matrix or even plain light microscopy images could be quantified using appropriate imaging techniques and software, Alizarin Red offers an additional way of quantification: the amount of dye binding to the mineral correlates to the amount of mineralization, after imaging the dye can be released from the minerals using acetic acid and can then be quantified using colorimetric spectrophotometry (27). Surface analysis was used in a similar way to study formation on scaffolds, films, or particles. Scaffolds were stained and/or imaged, and the area of matrix deposition was visualized or quantified. Volume analysis was used to describe the measurement of mineralized tissue components calcium and phosphate, which were released after destruction of the matrix. These three types of formation measurement are destructive methods, meaning that the samples must be sacrificed for each time point.

The remaining two types of formation methods are non-destructive. Supernatant analysis was used to describe the measurement of Collagen type I C-terminal propeptide (CICP), a byproduct of collagen deposition, in cell culture supernatant. 3D scan was used to describe the use of (in this case) μCT quantify the three-dimensional structure of mineralized matrix.

#### TRAP measurements as a surrogate marker of osteoclastic resorption

TRAP is a protein that has long been used as the predominant OC marker (28). Out of all OB-OC studies in Database 1, TRAP was studied 63 times in 61 publications by a dedicated assay (Table 4). TRAP can be measured intracellularly or excreted into the medium in two ways. Its enzymatic phosphatase activity can be measured directly, or the amount of TRAP molecules present can be quantified. TRAP release was studied both on cell lysate and on supernatant, and in some cases on both. The most frequently used method to study TRAP activity was using 4-nithophenylphosphate (pNPP), a substrate that is cleaved by phosphatases into phosphate and detectable yellow 4-nithophenol. Others used the fluorophore Naphthol ASBI-phosphate, which is converted into the fluorescent Naphthol-ASBI (29) and shows specificity for TRAP isoform 5b, making this method more specific for the detection of OC when compared to the measurement of TRAP enzyme in general (30). Naphtol ASMX phosphate (31) and an otherwise undisclosed diazonium salt function in a similar manner. Enzyme linked Immunosorbent Assay (ELISA) can be used to detect TRAP in a slightly different manner; by binding a detectable substrate directly to the TRAP enzyme instead of using the enzyme to produce detectable substrate. Relying on conjugated enzymes or fluorescence, these techniques should be more effective at low concentrations of TRAP because multiple conjugates could bind to a single TRAP molecule. Others used a kit to detect TRAP, but no description of the assay other than the manufacturer were given.

**Table 4:** TRAP measurement techniques and analyses. Each column in Table 4 signifies a different technique to measure TRAP. This table consists of two distinct sections. The first section shows the number of studies that used each technique, and whether these were used on (lysed) cells or on culture supernatant. If only a reference to other literature was provided, that instance was listed in the row ‘Reference only’, and when these details were not reported, that instance was listed in the row ‘Not reported’. Note that in a single study TRAP can be measured with the same technique on both cell lysate and culture supernatant, resulting in a higher count of occurrences than number of studies that analyzed TRAP. The second section shows with which method of analysis the TRAP content was measured. If only a reference to other literature was provided, that instance was listed in the row ‘Reference only’, and when these details were not reported, that instance was listed in the row ‘Not reported’. If one study measured TRAP on both cells and supernatant, then that study is represented twice in the second section. In all other cases, each study is represented once in each section.

| Type | pNPP | N-ASBI-P | N-ASM-X-P | ELISA | Diazonium salt | Undefined kit | Reference | Not reported | Total |
| --- | --- | --- | --- | --- | --- | --- | --- | --- | --- |
| Total | 33 | 5 | 1 | 9 | 1 | 9 | 4 | 1 | 63 |
| Lysed cells | 29 | 5 | 1 | 1 |  | 3 | 2 |  | 41 |
| Supernatant | 6 |  |  | 7 | 1 | 6 | 2 |  | 22 |
| Reference only |  |  |  | 1 |  |  |  |  | 1 |
| Not reported |  |  |  |  |  | 1 |  | 1 | 2 |
| Analysis | pNPP | N-ASBI-P | N-ASM-X-P | ELISA | Diazonium salt | Kit | Reference | Not reported | Total |
| absorbance | 33 |  | 1 | 8 |  | 6 | 2 | 1 | 51 |
| Fluorescence |  | 5 |  |  |  |  |  |  | 5 |
| Radiography |  |  |  |  |  |  |  |  | 0 |
| Reference only | 2 |  |  | 1 | 1 |  | 2 |  | 6 |
| Not reported |  |  |  |  |  | 4 |  |  | 4 |

#### ALP measurements as a surrogate marker of osteoblastic tissue formation

Alkaline phosphatase (ALP), a bone turnover marker that is commonly used to investigate OBs, was studied in 42 publications (Table 5). ALP is a phosphatase, that like TRAP in OCs can be found both within and on the OBs surface and can be excreted into culture medium soluble or via extracellular vesicles (32,33). The most frequently used method to measure ALP was to use the substrate pNPP, which is cleaved by ALP into phosphate and detectable yellow 4-nitrophenol. Enzyme Immuno Assays (EIA) and ELISAs are similar immunoenzymatic assays (34) that rely on an labelling ALP molecules with a detectable substrate or other enzymes. This is in contrast with the pNPP-based methods, where the ALP enzyme itself through its inherent enzymatic activity is responsible for generating the colored substance. An advantage of the EIA and ELISA methods is that these are generally more sensitive; multiple detectable molecules or enzymes can be bound to each ALP molecule. Others used a kit to measure ALP, but no description of the assay other than the manufacturer were given.

**Table 5:**
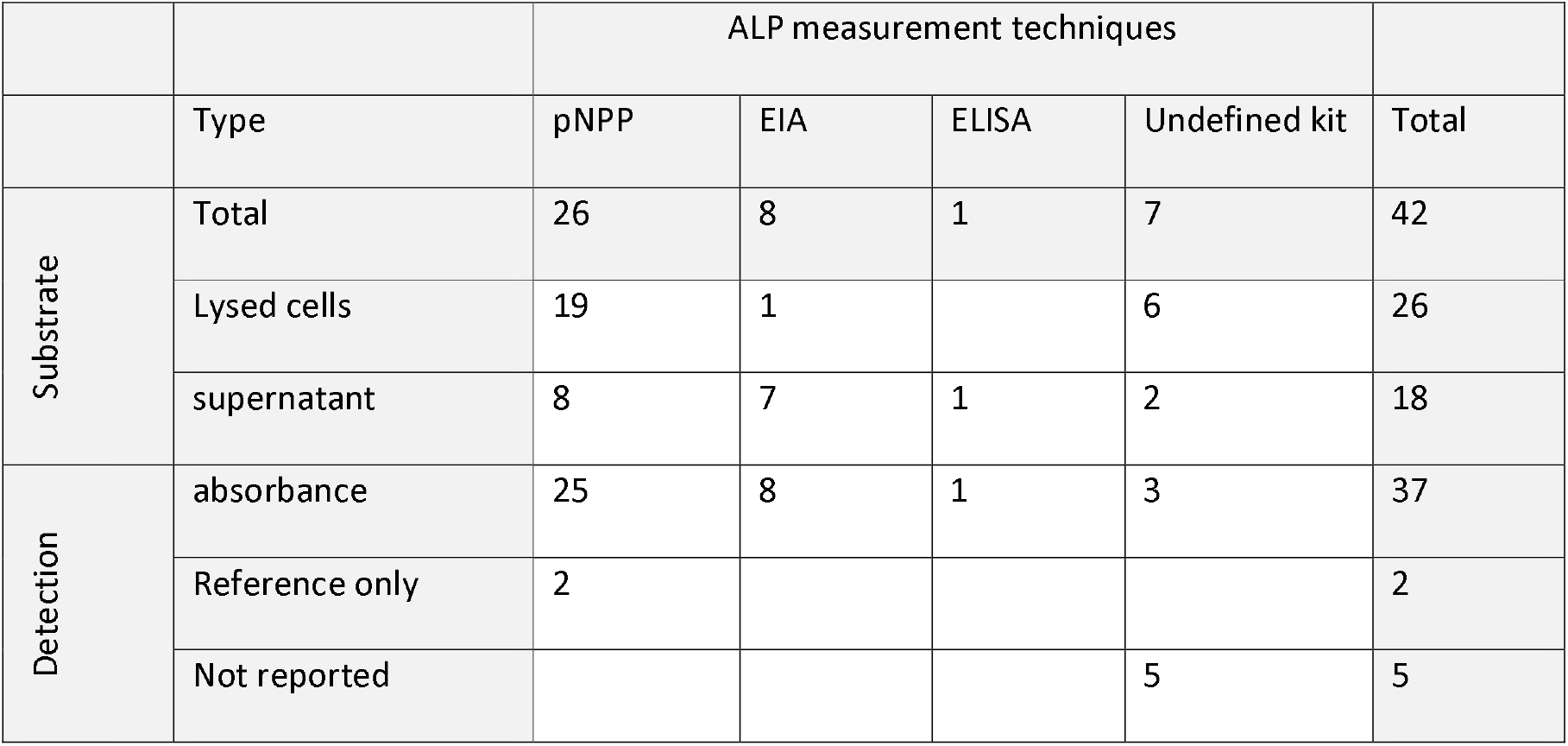
ALP measurement techniques and analysis. Each column signifies a different technique to measure ALP. The first rows show the occurrence of each technique and whether these were used on (lysed) cells, or on culture supernatant. Note that in a single study ALP can be measured with the same technique on both cell lysate and culture supernatant, resulting in a higher count of occurrences than number of studies that analyzed ALP. The final three rows show with which method of analysis the ALP content was measured.

### Database 2 results

While Database 1 was created to provide an overview of all reported methods to study the relevant outcome measures (resorption, formation, TRAP and ALP) without other experimental details, Database 2 was created to provide more insight into what culture conditions were used for cocultures. From Database 1, studies that investigated relevant outcome measures on both OB and OC were regarded as co-cultures capable of showing OB-OC interaction, versus using one cell type only to stimulate an effect or differentiation in the other. Of these qualifying studies, more information on the used cells and culture conditions was extracted and analyzed in Database 2.

#### Osteoblasts

Osteoblasts are the bone forming cells responsible for depositing mineralized matrix. From all 39 studies included in Database 2, the cell types that were present at the start of the co-culture were recorded and are shown in Table 6. More than half used human primary cells, whereas the others used animal primary cells or any type of cell line. Whether OBs or their progenitor cells were applied differed greatly between studies: almost half of the studies started the co-culture with OBs, the others started the co-culture with a type of progenitor cell. It needs to be noted that some of cell descriptions in Table 6 might refer to identical cell populations. This is a result of ambiguous isolation methods and nomenclature which is subjective and can evolve over time (35). This systematic map reflects the nomenclature used by the authors or an unambiguous translation of the provided nomenclature to nomenclature used in this map and does not interpret the provided information if it was ambiguous.

**Table 6:** Osteoblast origins and occurrences. From Database 2, the origin of the cells that were used as OB was extracted. Each column represents a different cell type of OB-like cells or their precursors. Each row represents a different source of cells, differentiating between both the origin species and whether the cells are primary cells or cell lines. Incremental totals are presented in the last row and column.

| Cell Origin | Osteoblasts | Mesenchymal stem cells | Mesenchymal stromal cells | Stromal cells | Stromal vascular Fraction | Osteoprogenitor cells | Per-row Total |
| --- | --- | --- | --- | --- | --- | --- | --- |
| Human primary | 4 | 9 | 2 | 6 | 1 |  | 22 |
| Human cell line | 1 |  |  |  |  |  | 1 |
| Mouse primary | 3 | 2 |  |  |  |  | 5 |
| Mouse cell line | 4 |  |  |  |  |  | 4 |
| Rat primary | 3 |  |  |  |  | 1 | 4 |
| Chicken primary |  |  |  | 2 |  |  | 2 |
| Reference only | 1 |  |  |  |  |  | 1 |
| Total | 16 | 11 | 2 | 8 | 1 | 1 | 39 |

One interesting observation regarding the cells used as OBs is that there is little variation in the different types of cells introduced into the co-cultures. Except for the oldest 6 studies that used chicken and rat cells, all studies used human or mouse cells, most of which were primary cells. While the studies using rat and mouse cells mostly directly introduced OBs (either isolated as such or differentiated before seeding), those that used human cells predominantly resorted to using progenitor cells (35). Such OB precursors can be obtained from blood and bone marrow donations and can be expanded to the required number of cells *in vitro.* The main difference between OB versus progenitors is the presence or absence of the osteogenic differentiation phase. Differentiation within the experiment could be desired for the research question or must be considered in case it is not. Those that used primary OBs purchased expandable human OBs (36) or used OBs (37), undefined expanded bone cells (38), or differentiated MSCs (39) from bone material obtained during a surgical procedure.

Seeding densities plays a major role in proliferation and cell function of OBs (18,83). Seeding density of OBs in 2D could be extracted or calculated for 26 studies and ranged from approximately 900 cells/cm^2^ to approximately 60.000 cells/cm^2^ with a median of approximately 6500 cells/cm^2^ and a mean of approximately 11000 cells/cm^2^ (Fig 4a). Seeding density of OBs in 3D could be extracted or calculated for 6 studies and ranged from approximately 300 cells/cm^3^ to approximately 7*10^7^ cells/cm^3^ with a median of approximately 4*10^6^ cells/cm^3^ and a mean of 15*10^6^ cells/cm^3^ (Fig 4d). It is important to note that these numbers are taken from the entire base of studies in Database 2, and as such are not representative for any type of OB or precursor used. These numbers can be further sorted and selected based on the individual researchers’ needs.

**Fig 4.**
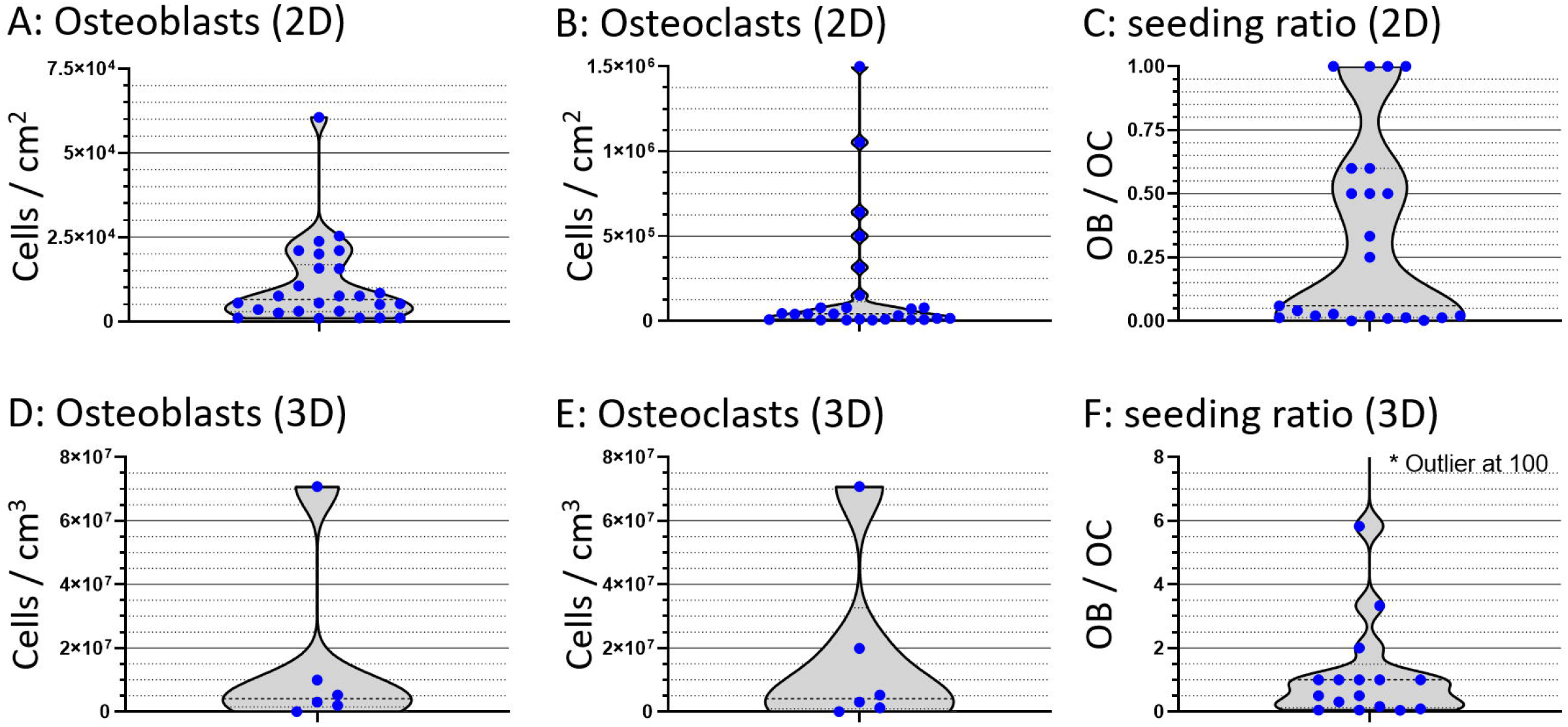
Seeding densities and seeding ratios. Violin plots of 2D and 3D seeding ratios of OB (A+D), OC (B+E) and respective seeding ratios (C+F). Values are calculated based on reported seeding numbers of the cells or precursors thereof by authors per surface are or volume. No distinction was made between different types of cells or precursors in these figures and this introduces a considerable spread in data due to possible cell proliferation (OB) and cell fusion (OC) that might have occurred after seeding. This distinction can be made in the database itself. Please take note that the ranges along the Y-axis are not the same for each figure. Each seeding density of each study is represented by a blue dot.

#### Osteoclasts

Osteoclasts are the bone resorbing cells that remove old and damaged bone tissue to make place for the deposition of new mineralized matrix. Out of all 39 studies included in Database 2, 20 used human primary cells, the others used animal primary cells or any type of cell line (Table 7). In most cases cultures were initiated with OC progenitors: 16 studies introduced monocytes, 11 introduced mononuclear cells, the rest used other precursors. Again, it needs to be noted that some of these descriptions are ambiguous. What is reported here is the definition used by the authors of the respective studies.

**Table 7:** Osteoclast origins and occurrences. From Database 2, the origin of the cells that were used as OC was extracted. Each column represents a different cell type of OC-like cells or their precursors. Each row represents a different source of cells, differentiating between both the origin species and whether the cells are primary cells or cell lines. If the cell source was indicated using only a reference, that instance was listed in the row ‘reference only’. Incremental totals are presented in the last row and column.

| Cell Origin | Monocytes | Mononuclear cells | Macrophages | Osteoclast precursors | Osteoclasts | Spleen cells | Total |
| --- | --- | --- | --- | --- | --- | --- | --- |
| Human primary | 10 | 6 | 1 | 3 |  |  | 20 |
| Human cell line | 4 |  |  |  |  |  | 4 |
| Mouse primary | 2 |  | 2 | 2 |  |  | 6 |
| Mouse cell line |  |  | 2 |  |  |  | 2 |
| Rat primary |  | 3 |  |  |  | 1 | 4 |
| Chicken primary |  | 2 |  |  |  |  | 2 |
| Reference only |  |  |  |  | 1 |  | 1 |
| Total | 16 | 11 | 5 | 5 | 1 | 1 | 39 |

The origin of the cells used as OCs is remarkably like those of the OBs. The 6 oldest included studies used chicken and rat cells, and all others used mouse or human cells. With only one exception combining a mouse ST-2 cell line with human monocytes (40), all studies used cells of exclusively a single species for the OB and OC source. Such a similarity was not found regarding the use of cell lines versus primary cells. While many studies introduced OBs directly into the co-culture, only a single study claimed to introduce OCs directly into co-culture but failed to provide any information regarding either cell source and was therefore ignored from further use.

Seeding density of OC in 2D could be extracted or calculated for 25 studies and ranged from 5*10^3^ cells/cm^2^ to 15*10^6^ cells/cm^2^ with a median of 42*10^3^ cells/cm^2^ and a mean of 190*10^3^ cells/cm^2^ (Fig 4b). Seeding density of OC in 3D could be extracted or calculated for 6 studies and ranged from 2*10^4^ cells/cm^3^ to 7*10^7^ cells/cm^3^ with a median of 4*10^6^ cells/cm^3^ and a mean of 17*10^6^ cells/cm^3^ (Fig 4e). Seeding ratios of OB:OC in 2D varied highly and ranged from 1:1500 to 1:1 (Fig 4c) and seeding ratios of OB:OC in 3D ranged from 100:1 to 1:25 (Fig 4f). In human bone tissue, the ratio of OB:OC is estimated to be approximately 7:1 (41). It must be noted that in these numbers, no distinction has been made between the use of precursors versus OB or OC or any type of expansion phase within experiments. These distinctions can be made within Database 2 for each individual need.

#### Co-culture medium composition and culture conditions

The behavior of cells is highly dependent on their environment, of which the biochemical part is predominantly determined by the culture medium composition. The main components of typical culture media are a base medium, fetal bovine serum (FBS) and specific supplements such as growth factors, especially when progenitor cells need to be differentiated first. Within the scope of this study, the base medium, FBS content and concentration of typical OB and OC supplements were analyzed. It became obvious that culture conditions are manifold and differ much between studies: A total of 8 different base (or complete) media were reported (Fig 5a), with αMEM and DMEM accounting for approximately 80% of all studies. FBS content ranged from 0% to 20%, with most studies using 10% (Fig 5b). Those without supplemented FBS used forms of complete media of which the composition was not described, but possibly including a type of serum or equivalent serum-free supplements.

**Fig 5.**
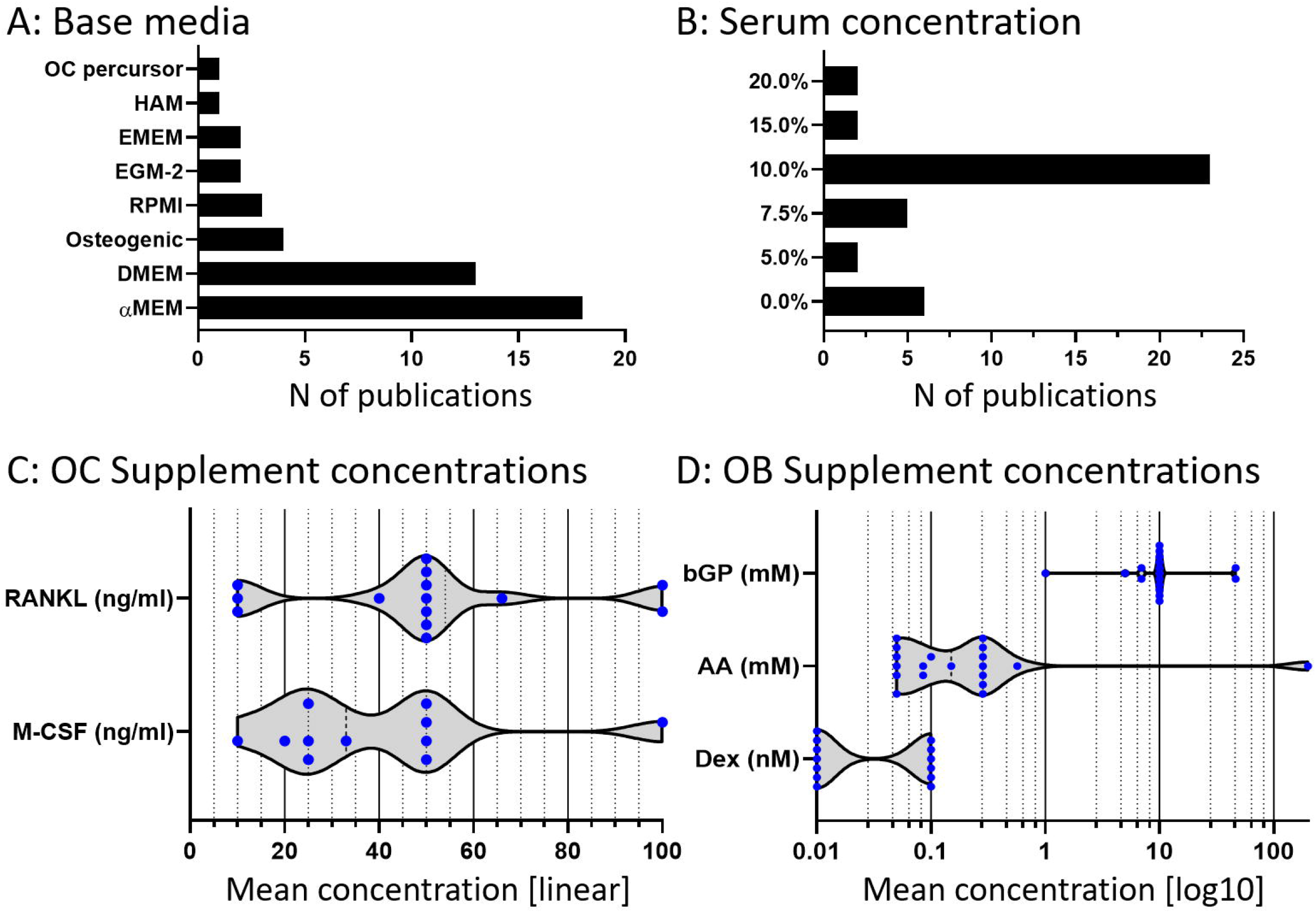
Medium components used by studies in Database 2. A) The occurrence of all identified base and complete media used during the co-culture phase of each study. B) Serum concentrations during the co-culture phase of each study. Numbers report exclusively the use of separately introduced FBS or FCS. Serum as part of a complete medium kit that was not described in the methods section is not reported here. C) OC supplements administered during the co-culture phase of each study. OC supplements were exclusively reported in ng/ml and are reported as such in the violin plot with all individual concentrations as blue dots. Please note that the x-axis has a linear distribution. D) Osteogenic supplements during the co-culture phase of each study. Osteogenic supplements were recalculated to molarity where necessary for comparability. Individual molarities are shown as blue dots. Please note that the x-axis has a logarithmic scale.

OC supplements were reported exclusively in ng/ml (Fig 5c). M-CSF concentration was reported in 11 studies and ranged from 10 ng/ml to 100 ng/ml with a mean of 39,82 ng/ml. RANKL concentration was reported in 14 studies and ranged from 10 ng/ml to 100 ng/ml with a mean of 49 ng/ml. All OB supplements were recalculated to molarity if they were reported in concentrations (Fig 5d). Ascorbic Acid (AA), which was also referred to as ascorbic acid-2-phosphate, L-ascorbic acid or L-ascorbate-2-phosphate, was used in 19 studies. AA concentration ranged from 0.05 mM to 0.57 mM, with mean of 0.18 mM and one outlier at 200 mM that was disregarded for this calculation. Dexamethasone was used and reported in 13 studies and was used in 2 different molarities: 6 times at 10^-7^ M and 7 times at 10^-8^ M. β-Glycerophosphate (βGP) use was reported in 17 studies, and ranged from 1 mM to 46 mM, with a mean of 13 mM.

#### Other culture conditions and techniques

In addition to the cell and medium characteristics, there are other factors that define an experiment. Out of the 39 studies of Database 2, 9 studies used a type of transwell or well insert culture (where cell populations are separated, but factor exchange is possible), 16 studies used a form of 3D culture, 3 studies reported the use of bioreactors, 2 studies used more than the required 2 cell types to form a tri- or tetra-culture (39,42), 7 studies reported using non-standard environmental conditions such as gas concentrations or mechanical loading. Polymerase chain reaction (PCR) was used in 13 studies, and supernatant analyses such as ELISA were used in 14 studies. The target genes, proteins or compounds were extracted from the publications and the occurrence of each target was recorded in the analysis of Database 2.

## Discussion

In recent years, many research groups have ventured into the realm of OB-OC co-cultures with the intent of studying both formation and resorption. Due to a lack of standardization within the field and the difficulty of finding publications based on methods instead of results, each group seems to be individually developing the tools to suit their needs resulting in many functionally related experiments that are methodologically completely different. The use of OB-OC co-cultures is usually not clearly mentioned in the title and abstract, making it difficult to find these studies without a systematic search and thorough review. The aim of this study was to generate a systematic map to give an overview of existing osteoblast-osteoclast co-culture studies published up to 6 January 2020, and present their methods, predetermined outcome measures and other useful parameters for analysis in 2 databases which can be filtered, sorted, searched and expanded.

The Database 1 contains all OB-OC co-culture studies in which at least one relevant primary outcome measure (formation and/or resorption) or secondary outcome measure (ALP and/or TRAP quantification as surrogate markers for formation and resorption, respectively) was investigated (S1_File_Database_1). A sub-selection of studies that have relevant outcome measures investigated on both OBs and OCs in the co-culture are shown in Database 2, accompanied by additional details on methods, culture conditions and cells (S2_File_Database_2).

### Resorption

Most studies in Database 1 investigating resorption did so in 2D cultures using a resorbable substrate such as bone, dentine, or synthetic osteological discs. This is not unexpected, as these three options are either the actual *in vivo* material (bone), a similar material with excellent properties for studying resorption (dentine) (43), or a material designed specifically for the purpose of studying resorption (osteologic discs or coated wells). One crucial advantage of using dentine discs over bone is related to the native structure of dentine itself: it does not contain canaliculi and has fewer other irregularities because it is not actively remodeled, providing more contrast between the native structure and resorption pits to accurately visualize them (43,44). Because of that reason, dentine is often favored over bone. The advantages of bone over dentine are that bone is the actual tissue of interest as opposed to a bone-like material, it can be obtained from many different species in relevant quantities and sizes, can be more easily be prelabeled *in vivo* with for example radioactive markers such as 3H-proline (45), it is cheaper and more readily available, and could be used in conjunction with cells from the same species or even same animal, although the latter was not observed in this map. Dentine is a component of ivory, usually obtained from elephants (46), hippo’s (47) or sperm whales (48). Regulations regarding ivory are strict and the material is rare, making it difficult and expensive to import and obtain. Synthetic osteologic discs have the advantage of being produced in a uniform manner and should show little sample-to-sample variation compared to discs made from animal tissue, or hand-made discs. Using well plates with thin osteologic coatings has the advantage that once the coating is resorbed, the translucent well below is revealed, which facilitates imaging with light microscopes. Combined with certain stainings, it makes quantifying resorbed area using conventional light microscopy easier.

Choosing the surface that will be resorbed by the osteoclasts will result in a compromise. For example, HA and other calcium phosphates are a likely choice for studying resorption since they are a major constituent of bone. While not optimized to facilitate resorption per se, they are simple to create, have a known composition and should offer good between-lab reproducibility. This contrasts with resorbable discs and plates with undisclosed ingredients and likely between-manufacturer variation. They are however synthetic, and do not contain any organic ECM components, which means that techniques such as measuring bone turnover markers NTx (49) and CTx (50) do not work.

It is believed that the deposition of collagen type I by osteoblasts is a vital step in the formation of mineralized tissue (51), and similarly could play a role in the resorption thereof. It is also possible to generate the to-be-resorbed material *in vitro* by the OBs (50), even within the same experiment. This essentially simulates a bone remodeling environment that is a step closer to the physiological process of bone remodeling versus only resorption, although *in vivo* the order in which this typically happens is reversed: first, damaged ECM is resorbed by OC, then new ECM is deposited by OB (52). However, the process of creating a mineralized matrix may introduce a variation in substrate size even prior to initiating the co-culture (53). Also, many *in vitro* formation experiments, while being able to produce the ECM constituents collagen and mineral, are not (yet) producing real bone ECM (51). An advantage specific to using a collagen-based material in favor of a pure ceramic material is that techniques such as NTx (49) and CTx (50) can be used. These bone turnover markers are used in the clinic and can quantify resorption by directly analyzing the liberated collagen fragments that were present in the resorbed mineralized matrix (54).

Because most studies were conducted in 2D, most resorted to using various types of 2D microscopy to analyze resorption, usually after staining to increase contrast. This can facilitate the quantification of resorbed area using image analysis software but is usually limited to a quantification of surface area, whereas resorption is a three-dimensional process. While methods exist to reconstruct a set of stereoscopic 2D images into 3D height maps (55), these were not identified within the studies in either database of this systematic map. It would be better to consider imaging techniques that can directly quantify the resorbed volume. Examples are 2-photon microscopy for thin samples and micro computed tomography (μCT) (56). Due to the non-destructive nature of μCT, it is well suited to monitor mineralized volume over time within the same samples (57) and images can be compared for changes over time (53,56). The usefulness of such a monitoring tool is however dependent on the envisaged resolution versus the corresponding potential cell-damage caused by radiation exposure (58,59). Registering consecutive images can even show both formation and resorption of mineralized tissue within the same set of images of the same sample if both mineralizing OBs and resorbing OCs are present (53). While μCT in this map is predominantly used on 3D samples, one study used it to quantify the thickness of mineralized films and combined that data with surface metrological data (60).

Overall, the golden standard (bone and dentine discs) remains the most-used method to study 2D resorption, although alternatives such as osteological coatings offer new and easy ways of quantification. Compared to 2D cultures however, 3D cultures are under-represented in this systematic map. While the systematic search covers all publications until January 6 2020 available, only 24 studies were labeled as 3D co-cultures in Database 1, the first being published only in 2006 (61). From these we learn that studying 3D resorption remains a challenge, with the only identified viable options for quantification being μCT imaging and supernatant analysis techniques such as NTx and CTx.

### Formation

The result of bone formation is the deposition of mineralized matrix. This is however a multi-step process of the presence of properly stimulated OBs that lay down a framework of type I collagen, which in turn is mineralized by the addition of calcium phosphates (51). No single method of measuring formation confirms the occurrence of each step in this process, instead relying on the assumption that the confirmed presence of one step indicates the presence of the entire process.

With most studies being two-dimensional co-cultures, it is no surprise that most formation analyses extracted from Database 1 were stainings. Of these, Alizarin Red is particularly interesting due to the option of quantifying the amount of bound dye, which correlates to the amount of calcium (27). A risk when using this method on larger samples is that it is not certain how far both dye application and dye extraction penetrate the material. This should not affect relative comparisons between different sample groups but could lead to underestimations of calcium deposition. By completely lysing the samples and directly measuring the exact amount of calcium or phosphate (62,63) this risk could be avoided, at the cost of not gaining information on the location and distribution of calcium or phosphate through the sample.

The two types of non-destructive formation measurements, CICP and μCT, are coincidently well-suited for the analysis of three-dimensional co-cultures as well. A major advantage of these is that because of their non-destructive nature, they can be used to measure the same samples over time, and they can be used prior to other destructive techniques. CICP measurements (64) have no negative effects on the co-culture, requiring only that culture supernatant samples can be taken at the desired timepoints, usually at medium exchange. The use of μCT leads to both quantification and visualization of mineralization within the same sample over time, but it has some aspects to consider. Most importantly, to use it as a non-destructive technique the samples must be cultured in sterile vessels capable of being scanned. This means that experiments are limited by severe practical constraints. Additionally, there is a direct correlation between the resolution of the images (and thus the minimal detectable size of mineral deposits) and exposure to radiation and subsequent cell damage (58,59). Radiation damage directly affects the usefulness as a monitoring tool, and a careful balance between minimal acceptable resolution and maximal radiation exposure must be found.

Overall, 2D nodule stainings were the most frequently used method to measure formation. Combined with Alizarin Red dye release these provide an easy way to quantify mineralization, though CICP supernatant analysis and μCT techniques provide a non-destructive alternative that can also be used for 3D co-cultures.

### ALP and TRAP

ALP and TRAP are the two major markers used for indirectly quantifying respectively OB and OC activity that were included into Database 1. Their presence is no conclusive proof that formation and resorption are occurring because ALP is expressed in differentiating MSCs already (65) and TRAP is expressed on monocytes as well (53), but there is a correlation between their presence and that of OB and OC, respectively. ALP is an enzyme that makes phosphates available to be incorporated into the matrix (66), while TRAP has been associated with migration and activation of OC (67). These enzymes can be measured both after lysis of the cells or within the culture supernatant. The former allows the quantification of enzyme per DNA content when combined with a DNA assay, whereas the latter allows the monitoring of relative enzyme release over time. The precise methodological details and experimental setup are of lesser importance for measuring ALP and TRAP than they are for measuring formation and resorption. All that is required is the possibility to use the supernatant or cell lysate, which is possible in most common experimental setups. The most frequently used methods are the pNPP-based methods where ALP and TRAP directly convert a substrate into a measurable compound. Napthtol-based methods (29) rely on a similar principle, and show an increased specificity for TRAP isoform 5B in particular (30). The main advantage of these methods is that they use the inherent enzymatic activity of ALP and TRAP, reducing the complexity and cost of the assay. However, the reliance on the inherent enzymatic activity of the enzymes is also a practical limitation as inherent activity can be affected by freezing and long-term storage. Especially when monitoring ALP or TRAP release over time, samples are commonly frozen and stored for different periods of time, and enzyme activity could be affected by this. A workaround would be to directly analyze the samples after taking them, or to use methods that rely on the presence and not the activity of these enzymes.

One of those methods is the immunoenzymatic assay, of which ELISA is the most well-known. With a traditional ELISA the antigen is first bound to the assay plate, and then labeled with one or a series of antibodies that are conjugated with an enzyme to convert a substrate to a chromogenic product (68). These methods have the capacity to detect lower concentrations of protein because it is possible to label each individual protein with an excess of new enzymes each capable of converting substrate. In the case of TRAP, ELISA kits exist that are specific for TRAP isoform 5b which is expressed almost exclusively in OCs (69), whereas isoform 5a is also expressed by macrophages and dendritic cells (70). While in a co-culture with pure populations of OB and OC this distinction would not be relevant, macrophages or macrophage-like cells can be used as a precursor for OCs (24), and thus express isoform 5a which could be detected in a pNPP based assay. Similarly, most co-cultures use a precursor or heterogeneous population that either contains macrophages or contains cells capable of differentiating into macrophages such as mononuclear cells (71), which means that the presence of other isoforms or even other phosphatases is likely. Whether this negatively affects the results is another matter that can only be determined by comparison between the two types of assay. Another factor to consider in co-cultures is the fact that both ALP and TRAP are phosphatases. Assays that rely on their inherent phosphatase activity may show cross-reactivity of other phosphatases, although this should be mitigated by controlling the pH during the test.

To conclude, pNPP based methods are the most frequently used methods for detecting ALP and TRAP due to their affordability and simplicity. However, immunoenzymatic detection methods are more sensitive and specific, and do not rely on the intrinsic enzymatic activity of ALP and TRAP which can be affected by freeze-thaw cycles, long-term storage, and could show cross-reactivity with other phosphatases.

### Osteoclasts

OCs are the bone resorbing cells, and together with bone forming OBs they keep the bone mass and bone strength in equilibrium with the required loads placed upon it. OCs are created when OC precursors such as monocytes exit the bloodstream because of chemotactic cues followed by the correct biochemical signals that result in cell-fusion into OCs. Cells are currently considered to be OCs when expressing TRAP, having an actin ring, and having at least 3 nuclei (6). Osteoclastic resorption *in vivo* is an integral part of bone maintenance. Old and damaged bone tissue is resorbed and quickly replaced by OBs with new bone tissue.

There is a clear preference in the studies identified for Database 2 for using human cells to generate OCs, most notably monocytes and mononuclear cells. These have in the past two decades proven to be a reliable and relatively straight-forward precursor population for OCs (6), they can be obtained from human blood donations, and are thought to be better representatives for studying human physiology than cells of animal origin (2,3).

The choice of using precursors versus differentiated OCs is forced sharply into one direction because of both biological and experimental limitations. The extraction of OCs from bone is possible but cumbersome, requires access to fresh bone material and generally does not yield relevant numbers of OCs. Generating OCs from circulating precursors has proven to be an easier way of obtaining OCs. However, OCs have an average life span of approximately 2 weeks (72,73), some of which would already be lost if OCs would be created prior to the actual experiments. In contrast to most cells, differentiation happens by fusion of several precursors into a single OC. Fused multi-nucleated OCs can become large and hard to handle without damaging them. For those reasons they are usually generated within the experiment itself instead of in a prior culture. In fact, the first OB-OC cocultures were designed specifically to generate OCs by using osteoblastic cell signals (9), as opposed to generating a model to study both OBs and OCs simultaneously as this systematic map has indexed (74).

OCs can currently be obtained *in vitro* without the need for OBs thanks to the discovery in 1999 that M-CSF and RANKL are the necessary and sufficient proteins to induce osteoclastic differentiation from precursors (10). The cells are predominantly introduced into the co-culture as precursors to differentiate within the co-culture, regardless of whether these two proteins are used or not. Where in the past researchers used spleen cells for this, the studies included in this systematic map predominantly use (blood-derived) mononuclear cells, monocytes, or macrophages as precursor cells. These four sources are closely related, and the main differences between them are the purity of the population and how far along the path to differentiated and active OCs they are. In short: Spleen cells contain many cells, among others mononuclear cells. A part of the mononuclear cell population consists of monocytes which are currently regarded as ‘the’ OC precursors (75,76). Monocytes can differentiate into macrophages or fuse together into OCs, depending on the biochemical cues received. Macrophage-like cell-lines are being used to generate OCs as well.

There are risks associated with each method of generating OCs. Animal cells introduce a between-species variation and can respond differently than human cells (17), human donor cells tend to exhibit large between-donor variation compared to cell lines (77) and the number of cells acquired is limited and variable (78). The large variation between donors again highlights the need for patientspecific disease models instead of generic bone models. By using cells of a single diseased donor, the reaction of that patient’s cells on potential treatment options can be studied. Immortalized cell-lines result in immortal subsequently generated OC-like cells. This is however not the case *in vivo* and while it can greatly reduce between-experiment and between-lab variation, it is also physiologically less relevant. While these risks and characteristics do not discredit any source as a viable source of OCs for any experiment, the results of the corresponding studies should be interpreted with these characteristics in mind.

### Osteoblasts

OBs are the bone forming cells, and together with bone resorbing OCs they keep the bone mass and bone strength in equilibrium with the required loads placed upon it. In addition to their role in bone formation, they excrete the exact biochemical cues necessary to generate OCs out of their circulating precursors. Before the identification and commercial synthesis of these factors, a coculture with OB was the only way to generate OCs *in vitro.*

The preference for the use of human primary cells identified in the studies included in Database 2 can be explained by the good availability of donor material, expandability of OB precursors, and because human cells have the potential to better reflect human physiology than cells from other species (2,3). The choice of OB progenitors versus OBs is not as crucial here as it is with OCs. MSCs, the most commonly used precursors, have a tri-lineage potential (79) and should be able to differentiate into OBs on a 1-1 ratio. The advantage of osteoprogenitors such as MSCs is that these are capable of extensive proliferation before differentiation and could be used to migrate into and populate hard-to-reach areas within 3D scaffolds. Additionally, using progenitors opens possibilities to study osteoblastogenesis in addition to bone formation. When the effect of an intervention on mineralization but not osteogenesis is under investigation, care must be taken that the intervention is not applied before differentiation is has been achieved.

The advantage of directly introducing OBs instead of precursors, whether obtained directly from primary material or pre-differentiated *in vitro,* is that these do not need to be differentiated within the experiment anymore, and all seeded cells are already OBs, and by extension, any experimental conditions affect only mature OBs and not osteoblastogenesis in parallel. Actual OBs or to-be-differentiated MSCs isolated from orthopedic surgery are the most common source of primary human OBs. However, healthy human donor OBs are scarce because they are mostly isolated after surgery of mainly diseased patients. Whether the use of OBs from unhealthy donors affects experimental results needs to be elucidated. On the other hand, using patient cells to create a personalized *in vitro* disease model is the first step towards personalized medicine, especially if all cells are of that same patient. Finally, the use of any type of animal cell instead of human cells carries the risk of finding inter-species differences that can affect the results and conclusions, and everything based on that, because animal cells can behave differently than human cells (17). While none of these risks directly discredit any of the methods obtaining OBs, the results must be interpreted with these risks and characteristics in mind.

### Culture conditions

The success of a cell-culture experiment is dependent on many factors related to culture conditions. For most cell-types, standard culture conditions have been established. During co-culture experiments however, the needs of two or more cell types need to be met. Medium components and factors may be needed in different concentrations, as they can be beneficial to one cell type but inhibitory to the other (80).

There is a clear preference for medium based on DMEM and αMEM, but the choice of base medium for a culture is not an easy one. Base media are generally chosen based on the intended cell type, recommendations by a manufacturer or supplier of either cells or medium, preferred effect on cells, interaction with other supplements, and earlier experience. These factors make direct comparison of experimental results by literature virtually impossible. Additionally, none of the studies mentioned why they specifically chose the base media they used.

Another variable in medium composition is the use of FBS (or FCS). It is commonly known that there can be batch-to-batch and between-brand differences in FBS (81) which can impact the results of an experiment tremendously. While different concentrations are being used, the most common FBS concentration is 10%. However, no study explains why each type and concentration of FBS was used.

Although there was no clear predictor for using or not using any of the osteoblastic or osteoclastic supplements, when they were used, the concentrations were usually within the same order of magnitude in all studies, except for ascorbic acid. However, only 2 studies used all 5 of the supplements indexed in this study (AA, βGP, Dexamethasone, M-CSF and RANKL) and many combinations of supplements have been registered in this map. Looking at OC supplements, it is generally accepted that RANKL and M-CSF are both necessary and sufficient for osteoclastogenesis (10). However, OBs can produce RANKL and M-CSF themselves to trigger differentiation (9) and therefore the supplements are not necessarily required in co-culture. The need for all osteoblastic supplements is not as great considering osteoblasts can be introduced in various stages of development. Still, each supplement contributes to a specific function. Dexamethasone upregulates osteogenic differentiation, βGP acts as a phosphate source, and AA is a co-factor involved in collagen synthesis (82). Depending on the type of cells introduced, the aim of the experiment and other methodological details, their inclusion could be beneficial.

Finally, many studies used or omitted specific supplements related to their research question regarding the activity of OBs or OCs or used less common supplements for differentiation such as vitamin D3, human serum or Phorbol 12-myristate 13-acetate. What is seldom addressed however, is the compromise that must be made in choosing the right supplements and concentrations. Adding too high doses of supplements could cause an excess of these signals in the culture medium, effectively overshadowing any other ongoing cell-signaling over the same pathway by other cells. This is of critical importance when the goal is not to achieve only OBs and/or OCs activity, but to obtain a homeostasis in which the two cell types regulate each other, with experimental conditions or interventions that are expected to affect this balance. Here, it may be beneficial to experiment with lower concentrations of factors, supplemented only during critical phases of the cells’ development or differentiation.

The choice of medium in a co-culture is most likely going to be a compromise and must be based on the exact research question to be addressed, where the advantages and disadvantages of base media and supplements for both cell types are carefully weighed. Most likely, the ultimate goal for the envisaged co-culture would be to reach tissue homeostasis, in which the environment is as similar to the *in vivo* environment in tissue homeostasis where cell interactions with each other can be monitored.

#### Seeding densities and seeding ratios

Using the correct seeding densities plays a major role in proliferation and cell function of OBs (18,83) and osteoclastic differentiation (84). The seeding densities reported in this map show an enormous spread. Many factors could have influenced these numbers. For example, some studies report the numbers prior to expansion, others expand the cells in (co-)culture. Similarly, the percentages of relevant precursor cells in heterogenous cell populations can vary widely. The cell numbers present and OB:OC ratio most likely even change during a co-culture due to ongoing cell-division, differentiation, fusion and different expected life spans and the corresponding cell death. Regrettably, the available documentation of exact cell numbers introduced is often lacking, and open to some interpretation. While the figures show this large spread in data points, the included databases can be manipulated to filter and select studies that match criteria according to the readers’ specific needs.

Animal type, cell type, cell line versus primary cells and even passage number may also directly influence the choice of seeding densities in addition to various experimental choices. At the same time, the purpose of the experiment and more specifically the purpose of the cells and type of interaction required should determine the necessary seeding density. Are the cells required to actively deposit or resorb measurable amounts of minerals, or are they just supposed to be there to facilitate OB-OC communication? The combination of all these factors suggests that there in fact is no one ideal seeding density, that the best density for a certain experiment can only be determined by taking all the above factors into account, learning from others that did a similar experiment, and most importantly verifying assumptions and predictions in the lab.

Looking at the cell seeding ratio, here reported as number of seeded OB/OB-precursors per seeded OC/OC-precursor, outliers can be normalized against their seeded counterparts. In 2D studies, there are never more OBs/OB-precursors than OCs/OC-precursors. At most, they are seeded at a 1:1 OB:OC ratio. Even though in human bone tissue the ratio of OB:OC is estimated to be approximately 7:1 (41), higher OC numbers than OB numbers are not unexpected. OB precursors can still proliferate, whereas OC precursors usually still need to fuse together to form mature OC or OC-like cells. In 3D we do not see the same trend, with ratio’s ranging from 1:20 to 100:1. These differences are again affected by the same factors that influence individual OB and OC seeding densities, further enhanced by the extra layer of complexity that are inherent to 3D cultures.

### Limitations of the systematic search

While the authors took great care to construct a series of search queries fine-tuned for each of the three online bibliographic literature sources, the authors cannot be certain that all relevant OB-OC co-cultures have been included into the two databases. The search was limited by the necessary addition of a ‘co-culture’ search element. Co-culture studies without any indication thereof in the title or abstract simply cannot be identified through the initial search. To compensate for this, screening step 4, searching through identified reviews and publications included into Database 2, was executed. The publications included into Database 1 or the complete list of identified OB-OC cocultures could have been screened for references as well, but the authors decided against this. Database 2 was specifically chosen for this because the likelihood of a publication that matches all relevant inclusion criteria citing other such publications was deemed high, whereas less relevant papers (included into Database 1, or not included at all) were considered much less likely to cite publications relevant to this systematic map that had not already been identified by the search itself or the screening of reviews and Database 2. Publications in languages other than English, Dutch or German were excluded because none of the researchers involved in data curation and analysis were fluent in those languages. No budget was available to hire a professional translator for the remaining languages. The consequence of that is that there is a likelihood that relevant publications were missed.

### Limitations of the databases

The use of co-culture models is a field that is still developing, and we are now aware that it is not only about adding an additional cell type, but that the complexity of such a culture is more than just doubled. The applied choice of methods, cells, and culture conditions should be tailored to the research question to be investigated, and ideally would be comparable to other studies within the field. This systematic map shows that the currently applied methods are far from standardized and that many research groups have developed their own approach attempting to overcome each challenge, making comparison between research groups virtually impossible. There is no consensus on cell types, seeding densities, seeding ratios or medium composition, and many of these are predominantly determined by the research question and whatever has been done before in each respective laboratory. For each study, 86 columns worth of data has been extracted including in some cases extrapolation and recalculation of numbers, which are now available for sorting and filtering for individual needs. Still these databases only scratch the surface of each study, and to fully understand the collected information and the context on which it was gathered, one must still read the full publication.

It must be noted that the quality of reporting in many cases is lacking. Both missing information critical and non-critical for reproducing the methods of the studies was identified, and only 13 out of 39 studies included in Database 2 did not miss at least a basic description of all indexed characteristics. However, more relevant details of these characteristics may have been omitted that describe exactly how each method or culture conditions was executed that were not required for this systematic map. Instead, this systematic map focuses on a high-level indexing and evaluation of defining characteristics of methods and culture conditions.

This systematic map is not intended to provide a definitive answer to the question of how to set up the perfect OB-OC co-culture. Instead, it allows searching through all relevant co-culture studies looking for specific matching experimental characteristics or culture details that may be applicable to one’s own research. For this, it contains the possibility to search, sort and filter through many relevant characteristics. This allows one to find relevant studies that may have already (partly) studied one’s research question, or that can be used as a guide to design comparable experiments.

## Conclusion

With this systematic map, we have generated an overview of existing OB-OC co-culture studies published until January 6, 2020, their methods, predetermined outcome measures (formation and resorption, and ALP and TRAP quantification as surrogate markers for formation and resorption, respectively), and other useful parameters for analysis. The two constructed databases are intended to allow researchers to quickly identify publications relevant to their specific needs, which otherwise would have not been easily available or findable. The presented high-level evaluation and discussion of the major extracted methodological details provides important background information and context, suggestions and considerations covering most of the used cell sources, culture conditions and methods of analysis. Finally, this map includes the instructions for others to expand and manipulate the databases to answer their own more specific research questions.

## Supporting information

Database 1

Database 2

Using the databases

List of all OB-OC co-cultures identified in initial ti-ab screening

PRISMA checklist

Systematic review protocol and Search Queries

## Supporting information

**S1 File. Database 1.** This database contains all studies in which at least one relevant outcome measure was investigated. Characteristics of outcome measures and descriptive statistics are listed in this database.

**S2 File. Database 2.** This database contains all studies in which at least one relevant outcome measure was investigated for both OB and OC. Characteristics of cells, methods and culture conditions, and descriptive statistics are listed in this database.

**S3 File. Using the databases.** This document provides instructions on how to operate the databases, how to add publications and expand the analyses with more elements.

**S4 File. List of all OB-OC co-cultures.** This list contains the initial list of 694 OB-OC cocultures obtained after screening, before full-text investigation and exclusion based on outcome measures.

**S5 File PRISMA checklist.** The PRISMA checklist describing all elements of the systematic review, and on what page or which section of the submitted manuscript to find them.

**S6 File Systematic Review Protocol and Search Queries.** The protocol and search queries as they were published prior to execution of the fulltext screening phase.

