## Supplementary material for "Osteoblast-osteoclast co-cultures: a systematic review and map of available literature": Using the databases

The two databases were constructed using Microsoft Excel. The scripts to analyze the databases were written in Excel Visual Basics (VBA), and are more commonly referred to as macro’s. To use the scripts, macro’s must be enabled and the files must be saved as ‘.xlsm’ macro-enabled documents. It is important to remember that the scripts work only on non-hidden data. Excel allows filtering the data based on numerous criteria, which makes not-selected data appear hidden. The hidden data is deliberately skipped, and all analyses and tables only contain data from the studies that were not hidden after filtering. Applying no filters means that all data is used. Filtering can be enabled from the ‘Data’ menu item, which then converts the 1^st^ row of the excel sheet into a header with all remaining rows containing the items to be sorted or filtered. Filtering should be enabled by default in both databases. The sort and filter functionality allow the user to quickly obtain relevant studies without having to make changes to the database. On the 3^rd^ sheet ‘Explanations’, a description is given for each row. To analyze a subset of (filtered) data, simply press the corresponding buttons present on the 2^nd^ sheet of the database called ‘Data’. The old data will be erased from the sheet, and new data will be written to the sheet.

Expanding the database

Just like any excel file, the databases can easily be expanded with new publications. However, for many columns the choice of words, order of words and even the use of capital letters are used specifically to allow the right information to be used by the script. Any deviation from this results in the script incorrectly handling the new publications. A good rule of thumb is to find an existing publication that contains similar data as the to-be-added publication and copy and adjust the data. The script should work on any number of publications, as long as the publications are included in the named table. To check this, go to the bottom right corner of the complete table. There should be a small triangle indicating the end of the table. Any publication included in the range from A1 to the cell containing that triangle is included in the analysis. Most words that are present on either the x or y axis of any of the tables on the ‘Data’ sheet are hard-coded into the script. If a searchable item is written not exactly as it is in the tables, it will most likely not be found by the script. This included the use of capital letters. If the item to be added does not fit into existing tables and the uses would like additional rows or columns in the table, the tables must be manually adjusted from within the script to include new searchable words or options. This requires some knowledge on VBA or coding in general.

Editing the scripts

The scripts used in the analysis are included with the databases as part of the ‘.xlsm’ file format. Each database contains independent scripts, only usable with that specific database. The scripts can be accessed using Excel developer options, which are hidden by default but can be enabled by adjusting the menu bar. Right click on an empty space in the menu bar, choose to edit and select the developer options to be added to the menu. Using the ‘comment’ functionality all scripts contain detailed explanations of what each line of code should do. Please take note that these scripts were written by a researcher with no formal education in coding or VBA who took on the challenge of figuring out how VBA works by himself. This was a learning process with a lot of trial and error, eventually resulting in the scripts as presented here. To assist with the correct wording and nomenclature, on the 3^rd^ sheet of each database (‘Explanations’), there is information on what each column should contain and how each script processes that information with tips on how to structure the information. Please note that first these 2 databases were constructed and only afterwards the scripts were written. To make the scripts work, wordings and descriptions were optimized for the script while working on the script. As a result, items that appear in both databases but are only analyzed in one of the two are not written and structured the same in both databases. Similarly, scripts cannot be simply copied between databases.
