## Supplementary material for "Osteoblast-osteoclast co-cultures: a systematic review and map of available literature": PRISMA checklist

### PRISMA 2009 Checklist

| Section/topic | # | Checklist item | Reported on page # |
|---|---|---|---|
| **TITLE** | | | |
| Title | 1 | Identify the report as a systematic review, meta-analysis, or both. First identified as systematic review. Later referred to as a systematic map. | 1, 4-5 |
| **ABSTRACT** | | | |
| Structured summary | 2 | Provide a structured summary including, as applicable: background; objectives; data sources; study eligibility criteria, participants, and interventions; study appraisal and synthesis methods; results; limitations; conclusions and implications of key findings; systematic review registration number. | 2 |
| **INTRODUCTION** | | | |
| Rationale | 3 | Describe the rationale for the review in the context of what is already known. | 4 |
| Objectives | 4 | Provide an explicit statement of questions being addressed with reference to participants, interventions, comparisons, outcomes, and study design (PICOS). Population: osteoblast-osteoclast co-cultures. Intervention: relevant techniques / outcome measures as described. Comparison: different techniques for same outcome measure. Outcome: A systematically structured map of characteristics of included studies, their methods and outcome measures. | 4-5 |
| **METHODS** | | | |
| Protocol and registration | 5 | Indicate if a review protocol exists, if and where it can be accessed (e.g., Web address), and, if available, provide registration information including registration number. Published via Zenodo, accessible through reference. | 5 |
| Eligibility criteria | 6 | Specify study characteristics (e.g., PICOS, length of follow-up) and report characteristics (e.g., years considered, language, publication status) used as criteria for eligibility, giving rationale. All criteria described in steps 1-4. | 6-11 |
| Information sources | 7 | Describe all information sources (e.g., databases with dates of coverage, contact with study authors to identify additional studies) in the search and date last searched. Pubmed, Embase and Web of Science on January 6, 2020. No authors were contacted. | 6 |
| Search | 8 | Present full electronic search strategy for at least one database, including any limits used, such that it could be repeated. All search strategies are publicly available through the reference to the review protocol. | 5 |
| Study selection | 9 | State the process for selecting studies (i.e., screening, eligibility, included in systematic review, and, if applicable, included in the meta-analysis). 4-step approach describes the entire selection process. | 6-11 |
| Data collection process | 10 | Describe method of data extraction from reports (e.g., piloted forms, independently, in duplicate) and any processes for obtaining and confirming data from investigators. All in duplicate, independently by at least 2 reviewers. | 7 |
| Data items | 11 | List and define all variables for which data were sought (e.g., PICOS, funding sources) and any assumptions and simplifications made. All variables and assumptions extensively defined in 4-step approach. | 6-11 |
| Risk of bias in individual studies | 12 | Describe methods used for assessing risk of bias of individual studies (including specification of whether this was done at the study or outcome level), and how this information is to be used in any data synthesis. Results of individual studies were not used. Instead, methods were examined and characteristics thereof were collected. | Not applicable |

| | | | |
|---|---|---|---|
| Summary measures | 13 | State the principal summary measures (e.g., risk ratio, difference in means). Collected relevant outcome measures plotted against other characteristics, and descriptive statistics thereof. Described in chapters 'Database 1 / 2 generation and analysis. | 11-15 |
| Synthesis of results | 14 | Describe the methods of handling data and combining results of studies, if done, including measures of consistency (e.g., $I^2$) for each meta-analysis. | Not applicable |



| Section/topic | # | Checklist item | Reported on page # |
|---|---|---|---|
| Risk of bias across studies | 15 | Specify any assessment of risk of bias that may affect the cumulative evidence (e.g., publication bias, selective reporting within studies). Selective reporting or omission of information is covered. No results are analyzed, nor is there a meta-analysis resulting in cumulative evidence. | 12-14 |
| Additional analyses | 16 | Describe methods of additional analyses (e.g., sensitivity or subgroup analyses, meta-regression), if done, indicating which were pre-specified. Database analysis with descriptive statistics. | 12, 14-15 |
| **RESULTS** | | | |
| Study selection | 17 | Give numbers of studies screened, assessed for eligibility, and included in the review, with reasons for exclusions at each stage, ideally with a flow diagram. Flow diagram is Fig. 1. Other data described in results. | Fig1, 15-16 |
| Study characteristics | 18 | For each study, present characteristics for which data were extracted (e.g., study size, PICOS, follow-up period) and provide the citations. Collected in Database 1 and Database 2, summaries presented in figures and tables of this study. | Database 1 + 2 |
| Risk of bias within studies | 19 | Present data on risk of bias of each study and, if available, any outcome level assessment (see item 12). Results not used for this systematic review/map. Not applicable. | N.A. |
| Results of individual studies | 20 | For all outcomes considered (benefits or harms), present, for each study: (a) simple summary data for each intervention group (b) effect estimates and confidence intervals, ideally with a forest plot. No results used for this systematic map, but characteristics of each study are presented in the databases. | Databases 1 + 2 |
| Synthesis of results | 21 | Present results of each meta-analysis done, including confidence intervals and measures of consistency. | N.A. |
| Risk of bias across studies | 22 | Present results of any assessment of risk of bias across studies (see Item 15). Selective reporting or omission of information is covered. Information is clearly presented in tables. | tables |
| Additional analysis | 23 | Give results of additional analyses, if done (e.g., sensitivity or subgroup analyses, meta-regression [see Item 16]). Databases contain all calculated results, figures and tables contain summary data. | Databases 1 + 2, figures and tables. |
| **DISCUSSION** | | | |
| Summary of evidence | 24 | Summarize the main findings including the strength of evidence for each main outcome; consider their relevance to key groups (e.g., healthcare providers, users, and policy makers). Each characteristic is extensively discussed. | 34-47 |

| | | | |
|---|---|---|---|
| Limitations | 25 | Discuss limitations at study and outcome level (e.g., risk of bias), and at review-level (e.g., incomplete retrieval of identified research, reporting bias). Two paragraphs are dedicated to limitations. | 47-49 |
| Conclusions | 26 | Provide a general interpretation of the results in the context of other evidence, and implications for future research. | 49-50 |
| **FUNDING** | | | |
| Funding | 27 | Describe sources of funding for the systematic review and other support (e.g., supply of data); role of funders for the systematic review. S.J.A.R. is supported by ZonMw More Knowledge with Fewer Animals Programme (MKMD), project number 114024141. S.J.A.R. and S.H. are financially supported by the European Union's Seventh Framework Programme (FP/2007-2013), Grant Agreement No. 336043 (project REMOTE). B.W.M.d.W., M.A.M.V. and S.H. are financially supported by the research program TTW with project number TTW 016.Vidi.188.021, which is (partly) financed by the Netherlands Organization for Scientific Research (NWO). | Funding section |

*From:* Moher D, Liberati A, Tetzlaff J, Altman DG, The PRISMA Group (2009). Preferred Reporting Items for Systematic Reviews and Meta-Analyses: The PRISMA Statement. PLoS Med 6(7): e1000097. doi:10.1371/journal.pmed1000097

For more information, visit: **www.prisma-statement.org**.
