## Supplementary material for "Osteoblast-osteoclast co-cultures: a systematic review and map of available literature": Systematic review protocol and Search Queries

### SYSTEMATIC REVIEW PROTOCOL FOR ANIMAL INTERVENTION STUDIES

FORMAT BY SYRCLE (WWW.SYRCLE.NL)
VERSION 2.0 (DECEMBER 2014)

| Item # | Section/Subsection/Item | Description | Check for approval |
|---|---|---|---|
| | A. General | | |
| 1. | Title of the review | Osteoblast-osteoclast co-culture models of bone-remodelling: A systematic review. | |
| 2. | Authors (names, affiliations, contributions) | **Stefan Remmers[1], Bregje de Wildt[1], Michelle Vis[1], Rob de Vries[2], Keita Ito[1,3], Sandra Hofmann[1]**<br>[1]Orthopaedic Biomechanics, Department of Biomedical Engineering and the Institute for Complex Molecular Systems,<br>Eindhoven University of Technology, Eindhoven, The Netherlands<br>[2]SYRCLE, Department for Health Evidence, Radboud Institute for Health Sciences, Radboudumc, Nijmegen, The Netherlands<br>[3]Department of Orthopaedics, UMC Utrecht, Utrecht, The Netherlands | |
| 3. | Other contributors (names, affiliations, contributions) | | |
| 4. | Contact person + e-mail address | Stefan Remmers, | |
| 5. | Funding sources/sponsors | European Union's Seventh Framework Programme (FP/2007-2013) / EU Project No. 336043 | |
| 6. | Conflicts of interest | | |
| 7. | Date and location of protocol registration | | |
| 8. | Registration number (if applicable) | | |
| 9. | Stage of review at time of registration | Selection stage | |
| | B. Objectives | | |
| | Background | | |
| 10. | What is already known about this disease/model/intervention? Why is it important to do this review? | Animal studies are morally problematic, expensive, time consuming and often not accurate or not suitable for translation to humans. Adaptation of suitable *in vitro* models could provide an additional tool of selection prior to animal experiments, effectively reducing the number of experiments and the number of experimental groups needed. Many groups, including our own, have recently attempted to develop an *in vitro* model of bone remodelling. Osteoblasts (bone forming cells) have been studied extensively and can be used reliably to show mineral deposition. The addition of osteoclasts (bone resorbing cells) often proves difficult, especially if quantifiable outcome measures are desired. All groups work on their own model system, and all use different methodologies and cell-culture conditions. Culture conditions and analyses vary widely and are not standardized. The goal of this systematic review is |  |

| | | |
|---|---|---|
| | | therefore to identify all relevant co-culture studies of osteoblast- and osteoclast-like cells, in order to determine which methods result in quantifiable outcomes and facilitate the use of quantifiable outcome measures to measure bone formation and resorption, with the aim of developing a standard for co-culture models of bone-remodelling. |
| | Research question | |
| 11. | Specify the disease/health problem of interest | The co-culture models can be used for fundamental research on bone remodelling, studying bone diseases such as osteoporosis, drug development, and personalized medicine. |
| 12. | Specify the population/species studied | Co-cultures of either human or animal primary cells or cell lines that contain both osteoblasts and osteoclasts |
| 13. | Specify the intervention/exposure | The cell-culture model exposed to plain osteoblastic and / or osteoclastic culture medium, supplemented with any kind of biochemical variation, or exposed to mechanical loading. |
| 14. | Specify the control population | The cell-culture model exposed to plain osteoblastic and / or osteoclastic culture medium |
| 15. | Specify the outcome measures | Measured osteoblast activity or bone formation (mineralization / calcium deposition), and measured osteoclast activity or bone resorption. |
| 16. | State your research question (based on items 11-15) | Main question: Which are the current osteoblast-osteoclast co-culture models?<br>Subquestions:<br>- Which of those have quantifiable outcome measures on bone formation or resorption?<br>- What methodological approaches and experimental conditions facilitate simultaneously studying changes in resorption and formation best? |
| | C. Methods | |
| | Search and study identification | |
| 17. | Identify literature databases to search (*e.g.* Pubmed, Embase, Web of science) | X MEDLINE via PubMed X Web of Science<br>☐SCOPUS X EMBASE<br>☐Other, namely:<br>☐Specific journal(s), namely: |
| 18. | Define electronic search strategies (*e.g.* use the step by step search guide[15] and animal search filters[20, 21]) | When available, please add a supplementary file containing your search strategy: [insert file name] |
| 19. | Identify other sources for study identification | X Reference lists of included studies ☐Books<br>X Reference lists of relevant reviews<br>☐Conference proceedings, namely:<br>☐Contacting authors/ organisations, namely:<br>☐Other, namely: |
| 20. | Define search strategy for these other sources | Of all studies included after the final screening phase and all related reviews, the titles in the reference lists will be |

| | | |
|---|---|---|
| 21. | | screened for additional includable studies. If from the title is seems likely that a study could match the criteria, it is added to a list of new studies, that are screened in the same way as the originally included studies. |
| | Study selection | |
| 21. | Define screening phases (*e.g.* pre-screening based on title/abstract, full text screening, both) | • Phase 1: Pre-screening based on title/abstract for bone-related co-culture primary studies<br>• Phase 2: full-text screening. In case of doubt in phase 1, screen full text for bone-related co-cultures.<br>• Phase 3 (within review): full-text screening for relevant outcome measures<br>• Phase 4 (within review): full-text screening for applicability as a model for bone remodeling |
| 22. | Specify (a) the number of reviewers per screening phase and (b) how discrepancies will be resolved | a. 2 reviewers will independently screen titles, abstracts and full texts.<br>b. If the reviewers disagree, they will discuss together. If no consensus can be reached, a 3rd reviewer will be consulted. |
| | *Define all inclusion and exclusion criteria based on:* | |
| 23. | Type of study (design) | Inclusion criteria: experiment is a primary study<br><br>Exclusion criteria: not a primary study. Reviews on the subject (will be used as an additional source to locate studies), conference abstracts |
| 24. | Type of animals/population (*e.g.* age, gender, disease model) | Inclusion criteria: Study uses osteoblastic and osteoclastic cells, or cell lines and/or progenitor cells of those cells, as long as they are expected to differentiate into the required cell types. A heterogeneous cell mix is allowed if it is related to the osteoblastic or osteoclastic lineages, and the fraction of necessary precursors of finally differentiated cells is measured.(e.g. blood derived mononuclear cells, mesenchymal stromal cells) The addition of a 3rd or 4th cell type is permitted, as long as the aforementioned 2 are present as well. Source may be human or any animal. Study is bone related.<br><br>Exclusion criteria: Does not contain osteoblasts AND osteoclasts, or related/progenitor cells. Study is not bone related. |
| 25. | Type of intervention (*e.g.* dosage, timing, frequency) | Intervention is not part of initial inclusion or exclusion criteria |
| 26. | Outcome measures | Inclusion criteria: Any outcome related to bone formation or resorption<br>Exclusion criteria: No bone-related outcome |
| 27. | Language restrictions | Inclusion criteria: No restrictions<br>Exclusion criteria: None |
| 28. | Publication date restrictions | Inclusion criteria: No restrictions<br>Exclusion criteria: None |
| 29. | Other | Inclusion criteria: All unique studies<br>Exclusion criteria: If deemed necessary, the same |

| | | |
|---|---|---|
| | | experiment lead to different publications. These publications will not be excluded, but if discovered will be grouped and treated as one large study. In a similar way, recurring methods by the same group will be treated as one. |
| 30. | Sort and prioritize your exclusion criteria per selection phase | Selection phase 1 (tiab):<br>- Not both osteoblasts and osteoclasts<br>- No in vitro study<br>- Not a primary study<br>- Not bone related<br><br>Selection phase 2 (full text):<br>Same as in phase 1 for those studies where title/abstract was not sufficient. The result of this is a comprehensive list of all osteoblast-osteoclast co-cultures.<br><br>Phase 3 (outcome measures; within review):<br>Within these results, all included studies will be further assessed on investigating the desired outcome measures using full text screening. Inclusion criteria in order of importance are as follows:<br>-Resorption as outcome measure<br>-Formation as outcome measure<br>-Quantified osteoblast (ALP) or osteoclast (TRAP) activity<br>Studies with other outcome measures or interesting methods that could be of interest are included on a separate list as well.<br><br>Phase 4 (applicability to study bone remodeling): Each study included in phase 3 is characterized based on the ability to serve as a co-culture model using the following categories; 'Only osteoclasts are studied', 'Only osteoblasts are studied', 'Both osteoblasts and osteoclasts are studied'. Studies where only one cell type is studied will be placed on a separate list.<br><br>All studies placed on separate lists are used to extract information on their methods and outcome measures only. All studies in the main dataset where both celltypes are studied AND at least resorption, formation or cell activity is measured will be used completely. |
| | Study characteristics to be extracted (for assessment of external validity, reporting quality) | |
| 31. | Study ID (*e.g.* authors, year) | Authors, year |
| 32. | Study design characteristics (*e.g.* experimental groups, number of animals) | 1. Type of co-culture<br>2. Experimental / intervention groups<br>3. Timeline of experiment<br>4. Sample size |
| 33. | Animal model characteristics (*e.g.* species, gender, disease induction) | 1. Type of osteoblastic cells<br>2. Type of osteoclastic cells<br>3. Cell-culture details: seeding density, moment of |

| | | |
|---|---|---|
| | | seeding, medium compositions |
| 34. | Intervention characteristics (*e.g.* intervention, timing, duration) | Intervention, concentration, duration, timing |
| 35. | Outcome measures | Method of measuring bone formation/resorption, with a special interest in any quantifiable osteoblast activity, mineral deposition or bone formation, and quantifiable osteoclast activity or bone resorption. |
| 36. | Other (*e.g.* drop-outs) | Sample dropout rate, if available |
| | Assessment risk of bias (internal validity) or study quality | |
| 37. | Specify (a) the number of reviewers assessing the risk of bias/study quality in each study and (b) how discrepancies will be resolved | a. 2<br>b. 2 reviewers will independently assess the risk of bias/study quality of each study. If the reviewers disagree, they will discuss together. If no consensus can be reached, a 3th reviewer will be consulted. |
| 38. | Define criteria to assess (a) the internal validity of included studies (*e.g.* selection, performance, detection and attrition bias) and/or (b) other study quality measures (*e.g.* reporting quality, power) | ☐By use of SYRCLE's Risk of Bias tool<br>☐By use of SYRCLE's Risk of Bias tool, adapted as follows:<br>☐By use of CAMARADES' study quality checklist, e.g [22]<br>☐By use of CAMARADES' study quality checklist, adapted as follows:<br>X Other criteria, namely: adapted version of the OHAT/NTP in vitro risk of bias tool + reporting of sample size calculation |
| | Collection of outcome data | |
| 39. | For each outcome measure, define the type of data to be extracted (*e.g.* continuous/dichotomous, unit of measurement) | Bone formation (expressed as e.g. volume, surface area, percentage, relative gene expression, protein release, enzyme activity). - continuous<br><br>Bone resorption (expressed as e.g. volume, surface area, percentage, relative gene expression, protein release, enzyme activity ) - continuous |
| 40. | Methods for data extraction/retrieval (*e.g.* first extraction from graphs using a digital screen ruler, then contacting authors) | First from text, then from graph, finally from authors. |
| 41. | Specify (a) the number of reviewers extracting data and (b) how discrepancies will be resolved | a. 2 reviewers will independently extract the data from the papers<br>b. If the reviewers disagree, they will discuss together. If no consensus can be reached, a 3th reviewer will be consulted. |
| | Data analysis/synthesis | |
| 42. | Specify (per outcome measure) how you are planning to combine/compare the data (*e.g.* descriptive summary, meta-analysis) | Descriptive summary of results (statistically significant change in bone formation/resorption or not), while comparing different (co-)culture conditions |
| 43. | Specify (per outcome measure) how it will be decided whether a meta-analysis will be performed | As we do not want to compare the effect sizes between different studies, a meta-analysis is not useful. |
| | *If a meta-analysis seems feasible/sensible, specify (for each outcome measure):* | |

| | | |
|---|---|---|
| 44. | The effect measure to be used (*e.g.* mean difference, standardized mean difference, risk ratio, odds ratio) | - |
| 45. | The statistical model of analysis (*e.g.* random or fixed effects model) | - |
| 46. | The statistical methods to assess heterogeneity (*e.g.* $I^2$, Q) | - |
| 47. | Which study characteristics will be examined as potential source of heterogeneity (subgroup analysis) | - |
| 48. | Any sensitivity analyses you propose to perform | - |
| 49. | Other details meta-analysis (*e.g.* correction for multiple testing, correction for multiple use of control group) | - |
| 50. | The method for assessment of publication bias | - |

Final approval by (names, affiliations): Date:

**Strategy**
**Search term groups:**
osteoblasts – osteoblast precursors – osteoclasts – osteoclast precursors – bone related words – co-culture related words

Search steps

Include studies mentioning osteoblasts OR osteoblast precursors in combination with bone related words
AND
Include studies mentioning osteoclasts OR osteoclast precursors in combination with bone related words
AND
include studies mentioning co-cultures

---

PUBMED

Osteoblast
Osteoblasts[mesh] OR Osteoblast*[tiab] OR Pre-osteoblast*[tiab] OR preosteoblast [tiab] OR osteoprogenitor*[tiab] OR ((Bone forming[tiab] OR osteogenic[tiab]) AND cells[tiab])

Osteoblast progenitor
Mesenchymal stem cells[mesh] OR Mesenchymal stem[tiab] OR Mesenchymal stromal[tiab] OR Mesenchymal progenitor[tiab] OR bone marrow stem[tiab] OR Bone marrow stromal[tiab] OR MSC*[tiab] OR Multipotent stem[tiab] OR Multipotent stromal[tiab] OR Multi-potent stem[tiab] OR Multi-potent stromal[tiab] OR Stromal cell*[tiab] OR Stromal fraction[tiab] OR Adipose derived stem[tiab] OR Adipose derived stromal[tiab] OR Adipose stem[tiab] OR Adipose stromal[tiab] OR Fat derived stem[tiab] OR Fat derived stromal[tiab] OR MC3T3*[tiab] OR Periodontal ligament cell*[tiab] OR MLO-Y4[tiab] OR MLO Y4[tiab] OR MLO A5[tiab] OR MLO-A5[tiab]

Bone related
bone resorption[mesh] OR Bone and Bones[mesh] OR Bone Development[mesh] OR osteoporosis[mesh] OR Osteopetrosis[mesh] OR bone formation[tiab] OR Resorption[tiab] OR resorbing[tiab] OR Bone remodeling[tiab] OR bone remodelling[tiab] OR Bone re-modeling[tiab] OR bone re-modelling[tiab] OR (Mineral*[tiab] AND matrix[tiab] AND deposit*[tiab]) OR Resorpt*[tiab] OR Osteoporosis[tiab] OR osteoporotic[tiab] OR osteopetrosis[tiab] OR osteopetrotic[tiab] OR bone diseas*[tiab] OR osteolysis[tiab]

Osteoclast
osteoclasts[mesh] OR osteoclast*[tiab] OR pre-osteoclast*[tiab] OR preosteoclast*[tiab] OR resorbing cells[tiab] OR odontoclast*[tiab] OR cementoclast*[tiab]

Osteoclast precursors
monocytes[mesh] OR Hematopoietic Stem Cells[mesh] OR Monocyte-Macrophage Precursor Cells[mesh] OR monocyt*[tiab] OR THP*[tiab] OR RAW264*[tiab] OR RAW 264*[tiab] OR RAW-264*[tiab] OR Mononuclear cell*[tiab] OR monoblast*[tiab] OR promonocyt* OR Mononuclear fraction[tiab] OR ((Hematopoietic[tiab] OR haematopoietic[tiab]) AND (stem[tiab] OR [stromal]))

Co-culture

coculture techniques[mesh] OR co-cult*[tiab] OR cocult*[tiab] OR coincubat*[tiab] OR co-incubat*[tiab] OR multi-cultur*[tiab] OR multicultur*[tiab] OR tri-cultur*[tiab] OR tricultur*[tiab] OR cultured together[tiab] OR cultivated together[tiab] OR Cell interaction*[tiab] OR Two cell types[tiab] OR (simultaneously[tiab] AND (seeded[tiab] OR cultured[tiab])) OR Bone model[tiab] OR Bone models[tiab] OR (Coupling[tiab] AND (bone OR formation OR resorption OR remodeling OR remodelling)) OR Resorption model[tiab] OR Model of resorption[tiab] OR Mixed culture[tiab] OR Synchronous cult*[tiab] OR Simultaneous cult*[tiab] OR binary cult*[tiab] OR dual culture[tiab] OR bi-cult*[tiab] OR colocalization[tiab] OR co-localization[tiab] OR colocalisation[tiab] OR co-localisation[tiab] OR heterocult*[tiab] OR heterotypic cult*[tiab] OR organoid[tiab] OR organotypic cult*[tiab] OR Trans-well[tiab] OR Transwell[tiab] OR Permeable barrier[tiab] OR Well insert[tiab] OR Well inserts[tiab] OR Well-insert[tiab] OR Well-inserts[tiab]

**Strategy**
**Search term groups:**
osteoblasts – osteoblast precursors – osteoclasts – osteoclast precursors – bone related words – co-culture related words

Search steps

Include studies mentioning osteoblasts OR osteoblast precursors in combination with bone related words
AND
Include studies mentioning osteoclasts OR osteoclast precursors in combination with bone related words
AND
include studies mentioning co-cultures

---

EMBASE

Osteoblast
exp osteoblast/ OR exp osteoblast cell line/ OR Osteoblast*.ti,ab,kw. OR Pre-osteoblast*.ti,ab,kw. OR preosteoblast.ti,ab,kw. OR osteoprogenitor*.ti,ab,kw. OR ((Bone forming OR osteogenic) AND cells).ti,ab,kw. OR (osteogen* AND cell*).ti,ab,kw.

Osteoblast progenitor
exp mesenchymal stem cell/ OR Mesenchymal stem.ti,ab,kw. OR Mesenchymal stromal.ti,ab,kw. OR Mesenchymal progenitor.ti,ab,kw. OR bone marrow stem.ti,ab,kw. OR Bone marrow stromal.ti,ab,kw. OR MSC*.ti,ab,kw. OR Multipotent stem.ti,ab,kw. OR Multipotent stromal.ti,ab,kw. OR Multi-potent stem.ti,ab,kw. OR Multi-potent stromal.ti,ab,kw. OR Stromal cell*.ti,ab,kw. OR Stromal fraction.ti,ab,kw. OR Adipose derived stem.ti,ab,kw. OR Adipose derived stromal.ti,ab,kw. OR Adipose stem.ti,ab,kw. OR Adipose stromal.ti,ab,kw. OR Fat derived stem.ti,ab,kw. OR Fat derived stromal.ti,ab,kw. OR MC3T3*.ti,ab,kw. OR Periodontal ligament cell*.ti,ab,kw. OR MLO-Y4.ti,ab,kw. OR MLO Y4.ti,ab,kw. OR MLO A5.ti,ab,kw. OR MLO-A5.ti,ab,kw.

Bone related
exp bone/ OR exp osteolysis/ OR exp bone development/ OR exp osteoporosis/ OR exp osteopetrosis/ OR bone formation.ti,ab,kw. OR Resorption.ti,ab,kw. OR resorbing.ti,ab,kw. OR Bone remodeling.ti,ab,kw. OR bone remodelling.ti,ab,kw. OR Bone re-modeling.ti,ab,kw. OR bone re-modelling.ti,ab,kw. OR (Mineral* AND matrix AND deposit*).ti,ab,kw. OR Resorpt*.ti,ab,kw. OR Osteoporosis.ti,ab,kw. OR osteoporotic.ti,ab,kw. OR osteopetrosis.ti,ab,kw. OR osteopetrotic.ti,ab,kw. OR bone diseas*.ti,ab,kw.

Osteoclast
exp osteoclast activity/ or exp osteoclast/ or exp bone marrow osteoclast cell/ OR osteoclast*.ti,ab,kw. OR pre-osteoclast*.ti,ab,kw. OR preosteoclast*.ti,ab,kw. OR resorbing cells.ti,ab,kw. OR odontoclast*.ti,ab,kw. OR cementoclast*.ti,ab,kw.

Osteoclast precursors
exp monocyte/ OR exp hematopoietic stem cell/ OR exp monocyte macrophage precursor cell/ OR monocyt*.ti,ab,kw. OR monoblast*.ti,ab,kw. OR promonocyt*.ti,ab,kw. OR THP*.ti,ab,kw. OR RAW264*.ti,ab,kw. OR RAW 264*.ti,ab,kw. OR RAW-264*.ti,ab,kw. OR Mononuclear cell*.ti,ab,kw.

OR Mononuclear fraction.ti,ab,kw. OR ((Hematopoietic OR haematopoietic) AND (stem OR stromal)).ti,ab,kw.

Co-culture

exp coculture/ OR exp organoid/ OR co-cult*.ti,ab,kw. OR cocult*.ti,ab,kw. OR coincubat*.ti,ab,kw. OR co-incubat*.ti,ab,kw. OR multi-cultur*.ti,ab,kw. OR multicultur*.ti,ab,kw. OR tri-cultur*.ti,ab,kw. OR tricultur*.ti,ab,kw. OR cultured together.ti,ab,kw. OR cultivated together.ti,ab,kw. OR Cell interaction*.ti,ab,kw. OR Two cell types.ti,ab,kw. OR (simultaneously AND (seeded OR cultured)).ti,ab,kw. OR Bone model.ti,ab,kw. OR Bone models.ti,ab,kw. OR (Coupling AND (bone OR formation OR resorption OR remodelling OR remodeling)).ti,ab,kw. OR Resorption model.ti,ab,kw. OR Model of resorption.ti,ab,kw. OR Mixed culture.ti,ab,kw. OR Synchronous cult*.ti,ab,kw. OR Simultaneous cult*.ti,ab,kw. OR binary cult*.ti,ab,kw. OR dual culture.ti,ab,kw. OR bi-cult*.ti,ab,kw. OR colocalization.ti,ab,kw. OR co-localization.ti,ab,kw. OR colocalisation.ti,ab,kw. OR co-localisation.ti,ab,kw. OR heterocult*.ti,ab,kw. OR heterotypic cult*.ti,ab,kw. OR organoid.ti,ab,kw. OR organotypic cult*.ti,ab,kw. OR Trans-well.ti,ab,kw. OR Transwell.ti,ab,kw. OR Permeable barrier.ti,ab,kw. OR Well insert.ti,ab,kw. OR Well inserts.ti,ab,kw. OR Well-insert.ti,ab,kw. OR Well-inserts.ti,ab,kw.

**Strategy**
**Search term groups:**
osteoblasts – osteoblast precursors – osteoclasts – osteoclast precursors – bone related words – co-culture related words

Search steps

Include studies mentioning osteoblasts OR osteoblast precursors in combination with bone related words
AND
Include studies mentioning osteoclasts OR osteoclast precursors in combination with bone related words
AND
include studies mentioning co-cultures

---

WEB OF SCIENCE

<u>Osteoblast</u>
TS=(Osteoblast* OR Pre-osteoblast* OR preosteoblast* OR osteoprogenitor* OR ((“Bone forming” OR osteogenic) NEAR/3 cells)

<u>Osteoblast progenitor</u>
TS=(“Mesenchymal stem” OR “Mesenchymal stromal” OR “Mesenchymal progenitor” OR “bone marrow stem” OR “Bone marrow stromal” OR MSC* OR “Multipotent stem” OR “Multipotent stromal” OR “Multi-potent stem” OR “Multi-potent stromal” OR “Stromal cell” OR “Stromal fraction” OR “Adipose derived stem” OR “Adipose derived stromal” OR “Adipose stem” OR “Adipose stromal” OR “Fat derived stem” OR “Fat derived stromal” OR MC3T3* OR “Periodontal ligament” NEAR/3 cell* OR “MLO-Y4” OR “MLO Y4” OR “MLO A5” OR “MLO-A5”)

<u>Bone related</u>
TS=(Bone AND (development OR formation OR tissue OR remodel* OR re-model* OR forming OR model) OR skelet* OR bone* OR resorbing OR (Mineral* AND matrix AND deposit*) OR Resorpt* OR Osteoporo* OR osteopetro* OR bone diseas* OR osteolysis OR ossification)

<u>Osteoclast</u>
TS=(osteoclast* OR pre-osteoclast* OR preosteoclast* OR “resorbing cells” OR odontoclast* OR cementoclast*)

<u>osteoclast precursors</u>
TS=(Macrophage NEAR/0 Precursor* OR macrophage-like NEAR/3 cell* OR monocyt* OR monoblast* OR promonocyt* OR THP* OR RAW264* OR RAW NEAR/0 264* OR RAW-264* OR Mononuclear NEAR/0 cell* OR “Mononuclear fraction” OR ((Hematopoietic OR haematopoietic) AND (stem OR stromal)))

<u>Co-culture</u>
TS=(co-cult* OR cocult* OR coincubat* OR co-incubat* OR multi-cultur* OR multicultur* OR tri-cultur* OR tricultur* OR “cultured together” OR “cultivated together” OR Cell NEAR/0 interaction* OR Two NEAR/3 cell NEAR/3 types OR simultaneously NEAR/3 seeded OR simultaneously NEAR/3 cultured OR Bone NEAR/0 model* OR (Coupling NEAR/5 (bone OR formation OR resorption OR remodeling OR remodelling)) OR Resorption NEAR/2 model OR Mixed NEAR/3 culture OR

Synchronous NEAR/3 cult* OR Simultaneous* NEAR/3 cult* OR binary NEAR/0 cult* OR dual NEAR/0 culture OR bi-cult* OR colocalization OR co-localization OR colocalisation OR co-localisation OR heterocult* OR heterotypic NEAR/3 cult* OR organoid OR organotypic NEAR/3 cult* OR Trans-well OR Transwell OR “Permeable barrier” OR “Well insert” OR “Well inserts” OR “Well-insert” OR “Well-inserts”)
